# Species Dependent Metabolism of a Covalent nsP2 Protease Inhibitor with *in Vivo* Anti-alphaviral Activity

**DOI:** 10.1101/2025.01.13.632788

**Authors:** Mohammad Anwar Hossain, Abigail K. Mayo, Anirban Ghoshal, Sharon A. Taft-Benz, Elizabeth J. Anderson, Noah L. Morales, Katia D. Pressey, Ava M. Vargason, Kim L. R. Brouwer, Nathaniel J. Moorman, Mark T. Heise, Timothy M. Willson

## Abstract

RA-0002034 (**1**) is a potent covalent inhibitor targeting the alphavirus nsP2 cysteine protease. The species-dependent pharmacokinetics and metabolism of **1** were investigated to evaluate its therapeutic potential. Pharmacokinetic profiling revealed rapid clearance in mice, predominantly mediated by glutathione *S*-transferase (GST)-catalyzed conjugation. This metabolic liability contrasted with slower clearance observed in human hepatocytes and preclinical species such as rats, dogs, and monkeys. Cross-species studies confirmed the dominance of GST-driven metabolism in mice, whereas oxidative pathways were more pronounced in dogs. Despite rapid systemic clearance, **1** achieved antiviral efficacy in mice, reducing CHIKV viral loads in multiple tissues. Initial estimations of human hepatic clearance and half-life extrapolated from animal data indicate that b.i.d. dosing of **1** will be possible to maintain concentrations sufficient for antiviral activity in humans. These cross-species pharmacokinetic and metabolism studies support the continued evaluation of **1** as a promising anti-alphaviral therapeutic.

**Graphical Abstract:** 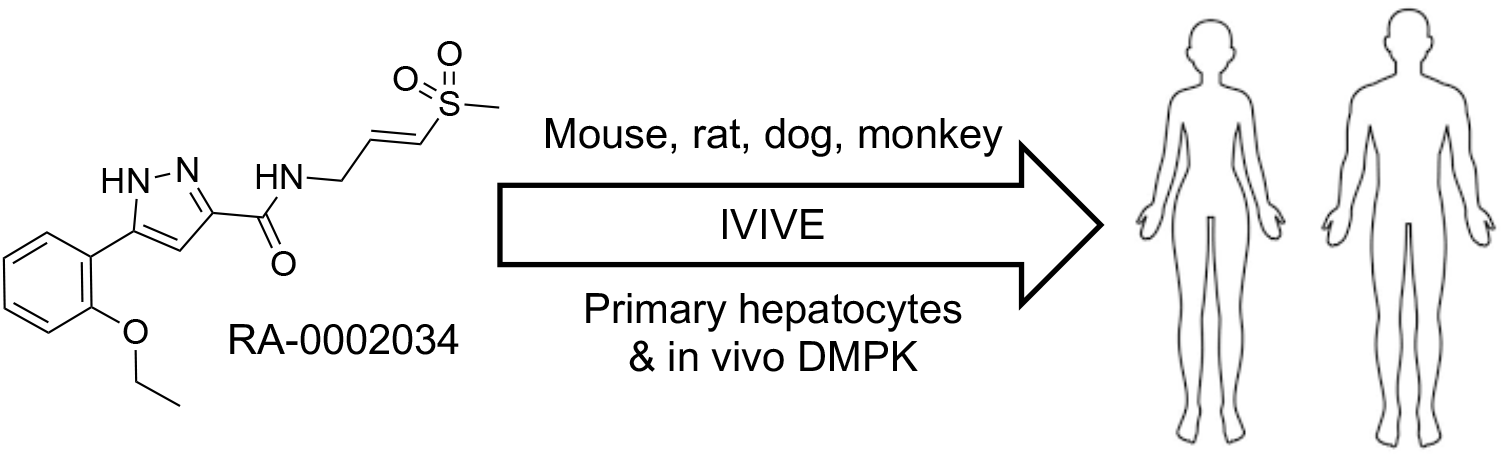

## Introduction

Covalent enzyme inhibitors have been developed as effective direct acting drugs to combat viral infections.^1^ Examples include nirmatrelvir that targets the main cysteine protease of SARS-CoV-2 and is used to treat COVID-19. Likewise, boceprevir and narlaprevir are covalent inhibitors of hepatitis C virus (HCV) NS3/4A serine protease and are effective in treatment of chronic HCV infections. These drugs show prolonged action due to covalent inactivation of their target enzymes. Modern chemoproteomic methods have allowed researchers to address one of major concerns of covalent enzyme inhibitor drugs, namely target selectivity across the proteome.^2^ However, one of the key remaining challenges is optimization of the pharmacokinetic properties of covalent drugs to support clinical advancement. On one hand, the chemical reactivity of electrophilic warheads makes them prone to conjugation and removal from the circulation by protective molecules such as glutathione resulting in short *in vivo* half-life values. This bioreactivity is countered by the fact that inactivation of the target protein may occur for several hours depending on resynthesis rate within the pharmacological system.^3^ Due to this potential disconnect between pharmacokinetics and pharmacodynamics, the optimization of drug exposure to maximize target engagement and inhibition with covalent antiviral drugs remains challenging.

RA-0002034 (**1**) is a selective nsP2 cysteine protease (nsP2pro) inhibitor with potent anti-alphaviral activity.^4, 5^ The vinyl sulfone warhead of **1** covalently captures the catalytic cysteine of nsP2pro and efficiently inactivates the enzyme with fast kinetics while showing remarkable proteome-wide selectivity.^5 6^ Anti-alphaviral activity was demonstrated in a *Chikunguny*a virus (CHIKV) replication assay, where **1** had an EC90 of 160 nM. **1** also produced a 5–8 log unit reduction of viral titer in infectious isolates of CHIKV, Venezuelan equine encephalitis virus, and Mayaro virus.^4, 5^ We previously reported that the acyclic **1** had a very short i.v. half-life in mice compared with its cyclic isomer **2**,^7^ suggesting that the vinyl sulfone moiety was a potential metabolic liability. In this report we describe the *in vivo* activity and species-dependent pharmacokinetics of **1**. Our studies indicate that mice may be unique in their ability to rapidly clear **1** from the systemic circulation. Importantly, cross-species profiling in primary hepatocytes and complementary *in vivo* pharmacokinetics support the continued preclinical development of **1** as a potential anti-alphaviral therapeutic drug.

## Results and Discussion

### ADME Properties and Mouse Pharmacokinetics

To support pharmacological testing of **1** the *in vivo* pharmacokinetics were initially determined in CD-1 mice by p.o. dosing at 30 mg/kg. The total plasma concentrations of **1** were quantified by LC-MS/MS (Table 1) and compared to the previously determined results from i.v. dosing at 10 mg/kg.^7^

**Table 1.** Mouse DMPK^a^.

|  | i.v. (10 mg/kg)<sup>b</sup> | p.o. (30 mg/kg) |
| --- | --- | --- |
| AUC<sub>last</sub> (h*ng/mL) | 875 | 860 |
| Vd<sub>ss</sub> (L/kg) | 1.04 |  |
| C<sub>max</sub> (ng/mL) |  | 1190 |
| t<sub>1/2</sub> (h) | 0.12 | 1.8 |
| CL<sub>s</sub> (mL/min/kg)<sup>c</sup> | 190 |  |
| F (%) |  | 32 |

<sup>a</sup>LC-MS analysis of pooled samples from CD-1 mice, n = 3. <sup>b</sup>Data from Ref. <sup>7</sup> <sup>c</sup>Based on plasma concentrations

Following i.v. dosing, **1** was rapidly cleared from the plasma compartment with a half-life <10 min, falling below the limit of quantitation within 3 h. Systemic clearance (CLs) was calculated as 190 mL/min/kg, equating to almost two times liver blood flow in a mouse and indicating a sizeable contribution of extrahepatic clearance to the elimination of **1**. Following p.o. dosing at 30 mg/kg (File S1), the plasma Cmax of **1** was 1190 ng/mL and it could be detected for 8 h. Using the i.v. and p.o. AUClast values, oral bioavailability was calculated as 32% (Table 1).

The *in vitro* metabolism of **1** was studied to identify the origin of its rapid systemic clearance. Although electrophilicity of the vinyl sulfone warhead in **1** was initially considered as a potential cause for concern, it was subsequently shown to be relatively unreactive to glutathione (GSH) conjugation with a half-life of 70 min in pH 7.4 buffer.^5, 7^ Likewise, when incubated in mouse whole blood, the half-life of **1** was >90 min (Table 2) suggesting that GSH conjugation in the plasma compartment was not the primary cause of its rapid clearance *in vivo*. Vinyl sulfone **1** also showed low clearance when incubated with mouse liver microsomes, with a half-life >60 min, indicating that hepatic P450 metabolism was likely to be only a minor contributor to its rapid systemic clearance.

**Table 2.** *In Vitro* Metabolism of 1^a^.

| | $t_{1/2}$<br>(min) | $CL_{\text{int}}$<br>($\mu\text{L}/\text{min}/10^6$ cells) |
| --- | --- | --- |
| Mouse whole blood | 97 |  |
| Mouse liver microsomes | 67 | 21<sup>b</sup> |
| Mouse hepatocytes | 13 | 105 |
| Human hepatocytes | 248 | 5.6 |
| Mouse hepatocytes + 1-ABT<sup>c</sup> | 26 | 54 |
| Mouse hepatocytes + EA<sup>d</sup> | 200 | 7.0 |

<sup>a</sup>5 $\mu\text{M}$ concentration. <sup>b</sup>Units of $\mu\text{L}/\text{min}/\text{mg}$ protein. <sup>c</sup>1 mM concentration. <sup>d</sup>0.2 mM concentration.

However, when **1** was incubated with primary mouse hepatocytes, which contain glutathione *S*-transferase (GST) enzymes not present in the microsomal fraction,^8^ it was very rapidly metabolized with a half-life of only 13 min and an intrinsic clearance of >100 μL/min/10^6^ cells (Table 2). Notably, metabolism of **1** was much slower in primary human hepatocytes. The results obtained in mouse hepatocytes suggested that GST-catalyzed metabolism may be a major contributor to the high clearance of **1** in mice *in vivo*. Co-dosing studies in primary mouse hepatocytes were performed to support this hypothesis. In the presence of 1-aminobenzotriazole (1-ABT), an irreversible inhibitor of P450 enzymes, the half-life of **1** was increased just 2-fold, confirming that phase I oxidation was only a minor component of its metabolism. In contrast, co-dosing with ethacrynic acid (EA), an irreversible inhibitor of GST enzymes,^9^ resulted in a dramatic 15-fold increase in the half-life of **1** indicating that GST-catalyzed GSH conjugation might be the primary driver of its rapid systemic clearance in mice.

To provide additional evidence that GST-catalyzed metabolism of **1** was responsible for its rapid clearance, GSH conjugation experiments were conducted in the presence of recombinant GST enzyme (Figure 2). GSH conjugation of **1** was efficiently catalyzed by the M1 isoform of mouse GST, reducing the half-life by 5-fold. Notably, the A1 isoform of human GST did not catalyze GSH conjugation, mirroring the species dependent metabolism observed in primary hepatocytes (Table 2). These results support the conclusion that **1** was a substrate for mouse GST, which catalyzed its GSH conjugation in the liver and other tissues resulting in a high systemic clearance. However, the lower intrinsic clearance of **1** in human hepatocytes and its stability to human GST-catalyzed GSH conjugation indicated that this metabolism may be species dependent.

**Figure 1.**
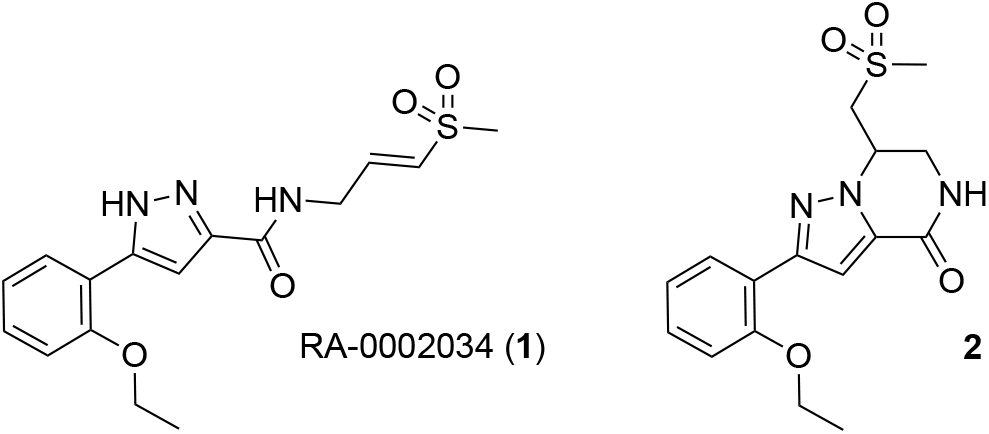
nsP2pro inhibitor RA-0002034 (**1**) and cyclic isomer **2**

**Figure 2.**
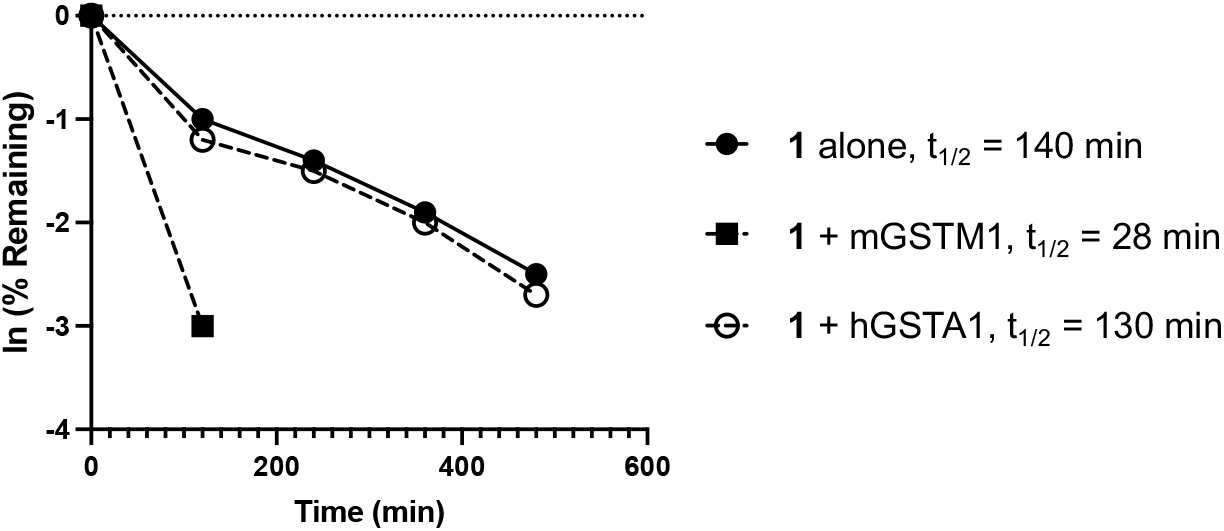
GST-catalyzed GSH conjugation of **1**.GSH (100x), phosphate buffer pH 7.4, mouse/human GST (10,000 ng/mL), 30 °C

### Metabolite Identification

Experiments in primary hepatocytes had indicated that **1** was a substrate for mouse P450 and GST enzymes. To corroborate that these pathways could account for the rapid clearance of **1** in mice, a study was performed to identify the metabolites formed *in vivo*. Pooled plasma samples collected over 5 h from mice dosed at 100 mg/kg p.o were analyzed by LC-MS/MS. A total of 18 metabolites were identified (Table 3 and Figure 3A) and their structures assigned by analysis of MS fragmentations (Figure 3B and S1).

**Table 3.** Metabolites of 1 in Mouse Plasma^**a**^.

| Peak | Time (min) | Mass (m/z) | -Alkyl | +GSH | + [O] | - [H] | [SO<sub>3</sub>] | Met | Glu | Hyd | +Acetyl | Assigned pathway | relative (%) |
| --- | --- | --- | --- | --- | --- | --- | --- | --- | --- | --- | --- | --- | --- |
| M1 | 4.78 | 326 | X |  | X |  | X |  |  |  |  | P450 | 1 |
| M2 | 5.53 | 498 | X |  |  |  |  |  | X |  |  | P450 | 35 |
| M3 | 5.60 | 514 | X |  | X |  |  |  | X |  |  |  |  |
| M4 | 5.62 | 444 |  |  | X |  | X |  |  |  |  |  |  |
| M5 | 5.72 | 542 |  |  | X |  |  |  | X |  |  | P450 | 4 |
| M6 | 6.06 | 590 |  |  | X |  |  | X |  |  |  |  | + |
| M7 | 6.16 | 546 | X | X |  |  |  | X | X | X |  |  | + |
| M8 | 6.29 | 526 |  |  |  |  |  |  | X |  |  | UGT | 4 |
| M9 | 6.50 | 606 |  | X | X |  |  | X | X | X |  |  | + |
| M10 | 6.70 | 657 |  | X |  |  |  |  |  |  |  | GST | 25 |
| M11 | 6.74 | 471 |  | X |  |  |  |  |  | X |  |  |  |
| M12 | 6.81 | 574 |  | X |  |  |  | X | X | X |  | GST | 16 |
| M13 | 7.02 | 462 | X |  |  | X |  |  | X | X |  |  | + |
| M14 | 7.49 | 232 | X |  |  |  |  |  |  |  |  |  | + |
| M15 | 7.75 | 513 |  | X |  |  |  |  |  | X | X |  | + |
| M16 | 8.06 | 233 |  |  |  |  |  |  |  | X |  |  | + |
| P | 8.15 | 350 |  |  |  |  |  |  |  |  |  |  | 15 |
| M17 | 8.34 | 350 |  | X | X | X |  | X |  | X |  |  | + |
| M18 | 8.97 | 398 |  | X |  |  |  | X |  | X |  |  | + |
| total (%) |  | - |  | 41 | 44 |  |  |  |  |  |  |  | 100 |

<sup>a</sup>See Figure 3B for structural assignments. Peak area percentages calculated from the UV absorption of each metabolite relative to parent drug in *t*<sub>0</sub> samples. '+': Only detected by MS. Met: Methylation, Glu: Glucuronidation, Hyd: Hydrolysis, UGT: UDP-glucuronosyltransferase.

**Figure 3.**
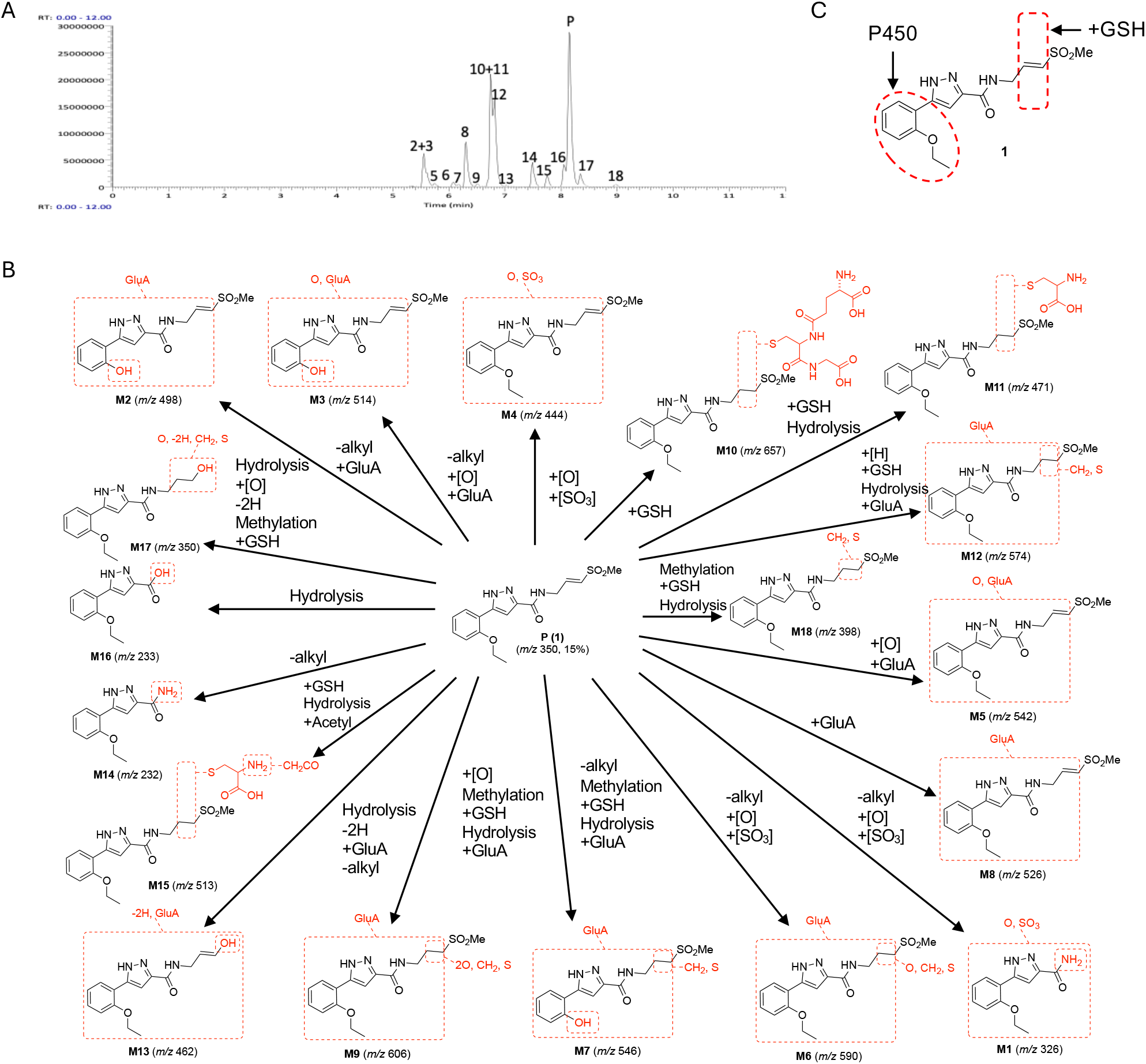
Metabolites of **1** in mice. (A) LC-MS of pooled plasma with identification of 18 metabolites of **1** (parent, P). X-axis represents total ion count. (B) Structural assignment of the metabolites (M1−M18) from MS fragmentation. Molecular ions are shown in parentheses. (C) Summary of the major sites of metabolism of **1** in mice.

Quantification by UV peak area of the metabolites (M1–18) relative to **1** (P) (Table 3) showed that that oxidation and GSH conjugation were the major pathways by which **1** was metabolized in mice (Figure 3C). The study indicated that the major circulating metabolites of **1** resulted from GSH conjugation of the vinyl sulfone (M10 and M12) or oxidation of the aryl substituent (M2) (Figure 3C), which was consistent with the analysis of *in vitro* metabolism in mouse hepatocytes. However, quantitation by UV peak area showed a ratio of ∼1:1 for metabolites formed by initial oxidation to those formed by GSH conjugation (Table 3). Additional minor metabolites resulting from oxidation of the λ-carbon vinyl sulfone sidechain (M1), epoxidation of the vinyl sulfone (M5), and glucuronidation of the pyrazole (M8) were also detected.

Additional validation was obtained through *in vivo* experiments using co-dosing of **1** in C57BL/6 mice with a P450 or a GST inhibitor (File S1). In C57BL/6 mice dosed with **1** (30 mg/kg) alone, drug could only be detected to the 5 h timepoint (Figure 4). Mice were pretreated with either EA (10 or 30 mg/kg p.o.) or 1-ABT (100 mg/kg p.o.) prior to dosing orally with **1**. At the 5 h timepoint (Figure 4), the plasma level of **1** was ∼10-fold higher and half-life was increased ∼2–3 fold in the mice co-dosed with either the GST or P450 inhibitor. The combined results of the primary hepatocyte, mouse pharmacokinetics, and metabolite identification studies indicated that **1** was substrate for P450 and GST enzymes and that a combination of phase I and phase II metabolism was responsible for its rapid *in vivo* clearance in mice (Figure 3C).

**Figure 4.**
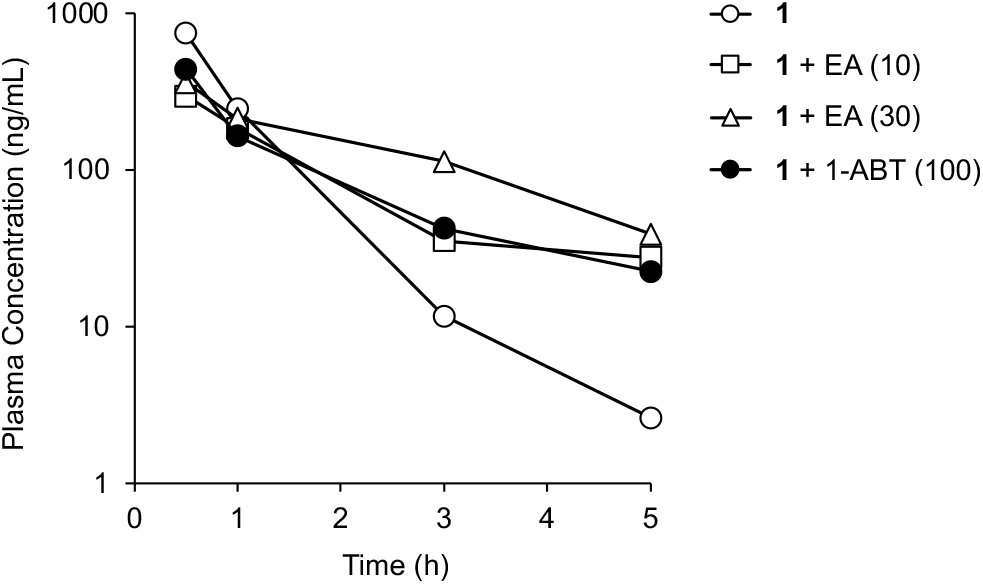
Clearance of **1** in mice. Male C57BL/6 mice dosed with **1** (30 mg/kg p.o.) in the presence of EA (10 or 30 mg/kg p.o.) or 1-ABT (100 mg/kg p.o.). EA was dosed twice at -6 and -2 h. 1-ABT was dosed once at -2 h. All data are the average from three mice.

### *In Vivo* Antiviral Efficacy

Although the clearance of **1** was rapid following i.v. dosing in mice, oral administration resulted in parent drug concentrations that could still be detected in the plasma of C57BL/6 mice at 5 h (Figure 4). To determine if sufficient levels of **1** could be sustained for 24 h in mice to potentially inhibit alphavirus replication an experiment was conducted using t.i.d. oral dosing of 100 mg/kg at 0, 8, and 16 h (Figure 5A and File S1). Plasma collected throughout the 24 h period indicated that total drug concentrations of **1** were sustained above the antiviral EC90 determined in the CHIKV-nLuc replication assay (160 nM = 56 ng/mL).^4, 5^ Measurement of mouse plasma protein binding determined that the free fraction (fu) of **1** was 18% of the total drug (Table S1). Adjustment of the plasma concentrations of **1** showed that while the unbound trough concentration was ∼10-fold below the EC90 for antiviral activity the peak was at or above this concentration. Since, **1** is an irreversible inhibitor of nsP2pro with fast kinetics (*Ki*_*nact*_/*Ki* > 6000),^5^ its ability to inactivate the viral enzyme *in vivo* may theoretically depend on the peak unbound drug concentration rather than total exposure.^10^ Therefore, the plasma concentration of **1** obtained with t.i.d. dosing at 100 mg/kg was deemed sufficient to attempt *in vivo* testing for antiviral activity.

**Figure 5.**
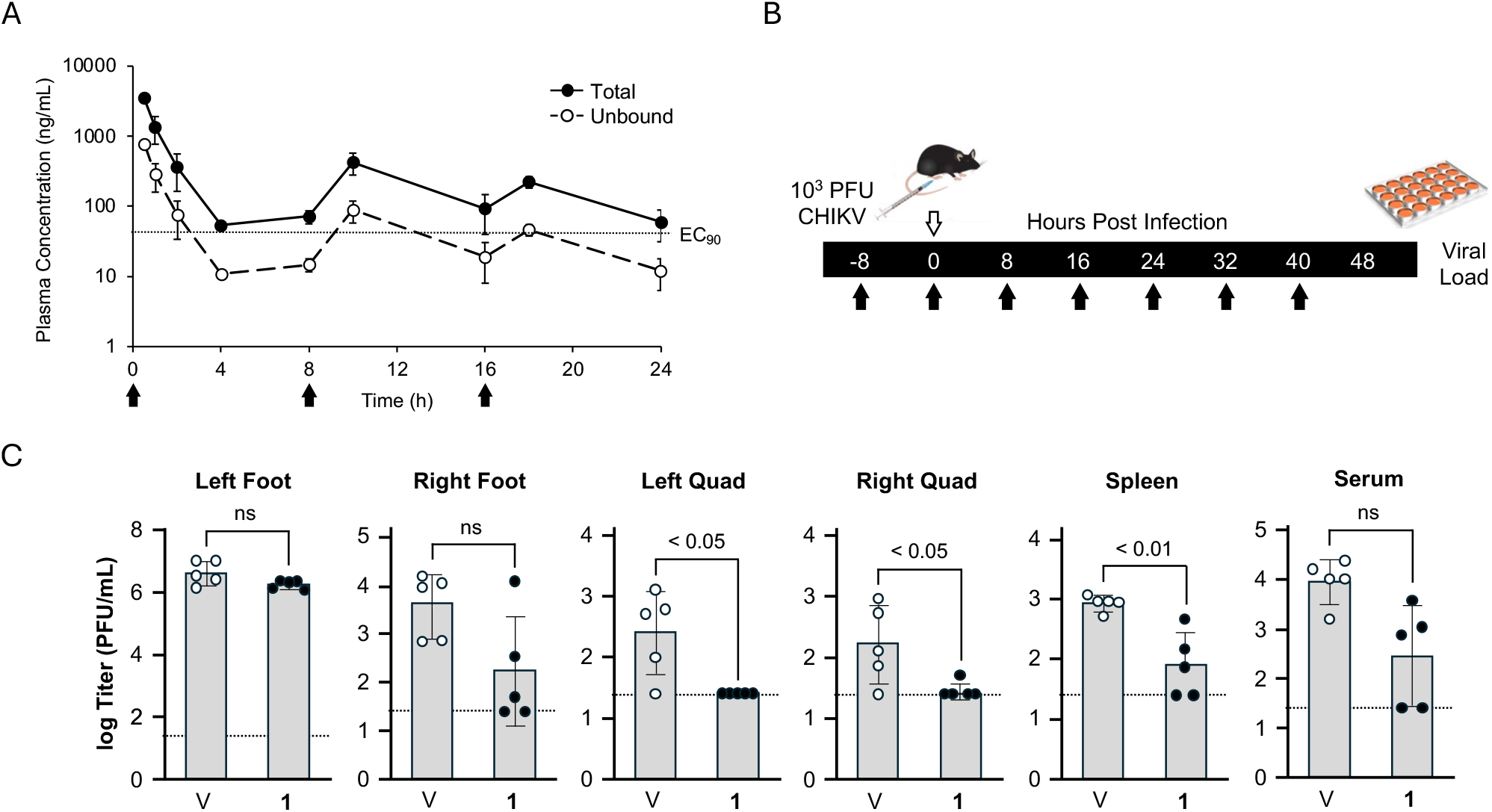
*In vivo* antiviral efficacy of **1**.(A) Plasma concentration of **1** in C57BL/6 mice (n = 3) dosed p.o. at 100 mg/kg t.i.d. Black arrows indicate the time of dosing. Solid circles are the total drug level. Open circles are the calculated unbound concentrations. The dotted line indicates the EC90 for inhibition of CHIKV replication (56 ng/mL). (B) Protocol for the mouse efficacy study. Black arrows indicate the time of dosing of **1** in mice (n = 5) at 100 mg/kg p.o. The open arrow indicates the time of inoculation with CHIKV (10^3^ PFU) into the left foot. Viral load was determined 8 h after the last dose. (C) Virus levels in 5 tissues and serum for individual mice (symbols) and group average (grey bars). The dotted line indicates the lower limit of quantification. Significance between **1** and vehicle (V) treated mice by t-test is indicated. ns = non-significant.

To test for antiviral efficacy, C57BL/6 mice were dosed orally at 100 mg/kg t.i.d. for 48 h (Figure 5B). Immediately prior to the second dose, the mice were inoculated with 10^3^ PFU of CHIKV in the left foot. Eight hours after the last dose, measurement of viral load in 5 tissues indicated that the levels were significantly elevated and were highest at the site of inoculation (Figure 5C). In the mice dosed with **1** no significant decrease in viral load was observed at the site of inoculation in the left foot. In the contralateral (right) foot, virus levels were reduced in 4/5 mice, but the group mean did not reach significance. However, in the left quad, right quad, and spleen significant decreases in viral load were measured. In the serum, a small but non-significant decrease in the circulating virus level was also observed (Figure 5C). The mouse *in vivo* study demonstrated that **1** had antiviral efficacy in multiple tissues even though it was not able to overcome the highest virus levels at the primary site of inoculation. *In vivo* testing of another covalent CHIKV nsP2pro inhibitor, J13, with similarly high systemic clearance in mice was recently reported.^11^ J13 also failed to decrease CHIKV viral titer at the site of inoculation, but its effect on virus levels in other tissues was not reported.^11^

### Cross-species Metabolism and Pharmacokinetics

In contrast to its rapid clearance in mouse hepatocytes, **1** was relatively stable in human hepatocytes (Table 2). To explore the species dependence of hepatic metabolism the stability of **1** was compared in cryopreserved primary hepatocytes from mice, rats, dogs, monkeys, and humans (Table 4).

**Table 4.**
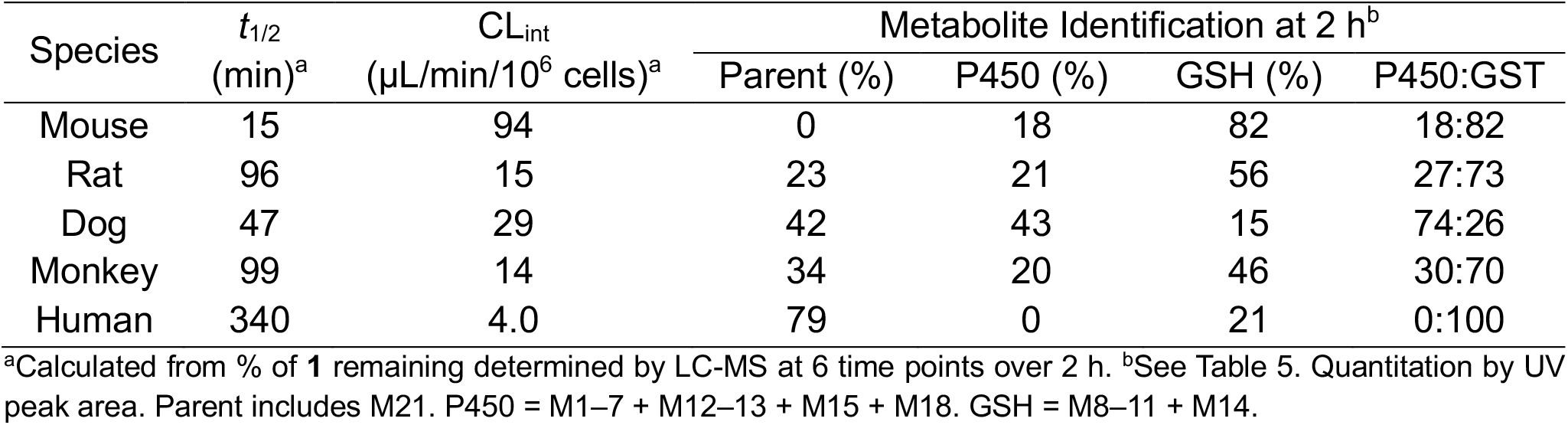
Cross-species Metabolism of 1 in Primary Hepatocytes.

| Species | $t_{1/2}$<br>(min)<sup>a</sup> | CL<sub>int</sub><br>(μL/min/$10^6$ cells)<sup>a</sup> | Metabolite Identification at 2 h<sup>b</sup> | | | |
| --- | --- | --- | --- | --- | --- | --- |
|  |  |  | Parent (%) | P450 (%) | GSH (%) | P450:GST |
| Mouse | 15 | 94 | 0 | 18 | 82 | 18:82 |
| Rat | 96 | 15 | 23 | 21 | 56 | 27:73 |
| Dog | 47 | 29 | 42 | 43 | 15 | 74:26 |
| Monkey | 99 | 14 | 34 | 20 | 46 | 30:70 |
| Human | 340 | 4.0 | 79 | 0 | 21 | 0:100 |

<sup>a</sup>Calculated from % of **1** remaining determined by LC-MS at 6 time points over 2 h. <sup>b</sup>See Table 5. Quantitation by UV peak area. Parent includes M21. P450 = M1–7 + M12–13 + M15 + M18. GSH = M8–11 + M14.

The highest rate of metabolism was observed in mouse hepatocytes and the slowest rate in human hepatocytes (Table 4), with intrinsic clearance values that closely matched the prior study (Table 2). The rate of metabolism measured in primary hepatocytes from the other three species was between the two extremes. Overall, the rank order of intrinsic clearance of **1** in primary hepatocytes was mouse > dog > rat ∼ monkey > human.

To identify the specific pathways of metabolism of **1**, samples were collected after 2 h of incubation with primary hepatocytes from each species and analysis performed by LC-MS/MS. Twenty-two metabolites were identified across the samples, with structures assigned by analysis of their MS fragmentation and quantitation from the UV peak area (Figure 6 and Table 5). In all species except mouse, a significant amount (23–79 %) of **1** (P) or its cyclized isomer **2** (M21)^7^ remained at 2 h. Glutathione conjugation as found in M8–11 was the major pathway of metabolism in mouse, rat, monkey, and human hepatocytes (Table 4). Notably, in human hepatocytes only GSH adducts were detected as major metabolites, with any P450 metabolites only detected by MS at levels too low to be quantified by UV peak area. However, in dog hepatocytes, oxidation (M12–M13) was seen as the major pathway of metabolism compared to glutathione conjugation (Table 4). Another notable pathway of metabolism in dog hepatocytes was dealkylation to the primary amide (M1–3, M7, and M16), which was most likely the result of P450 oxidation of the λ-carbon in the vinyl sulfone sidechain.

**Table 5.**
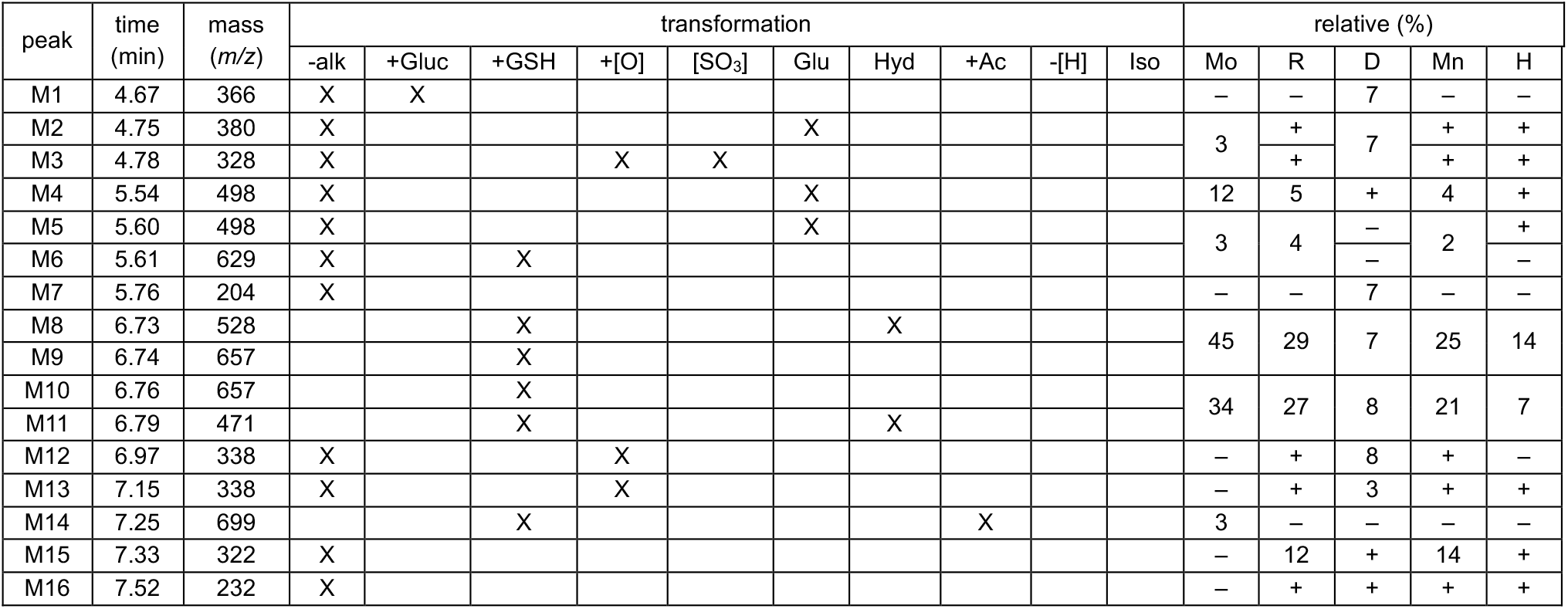
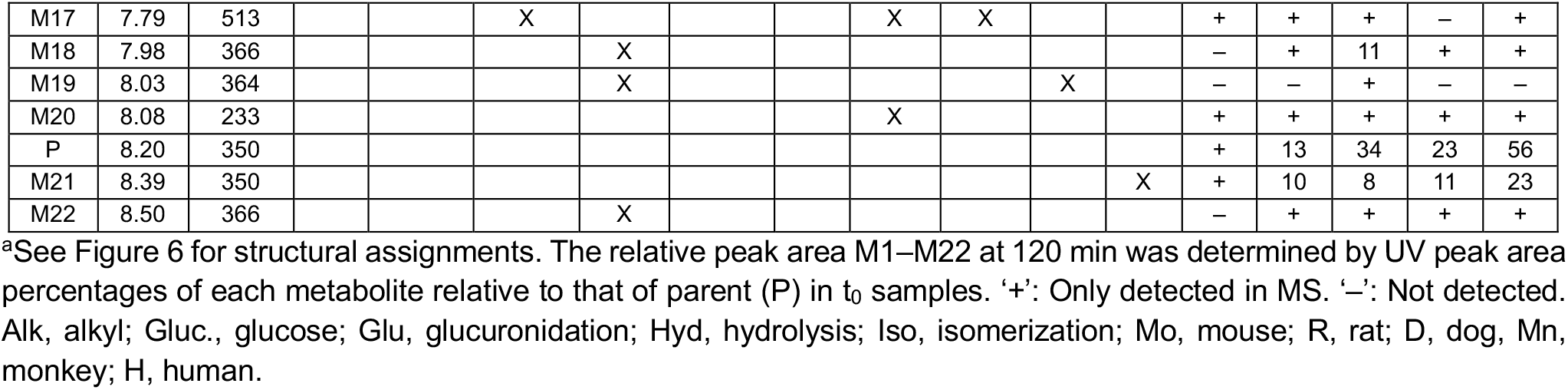
Cross-species Metabolites of 1^a^.

**Figure 6.**
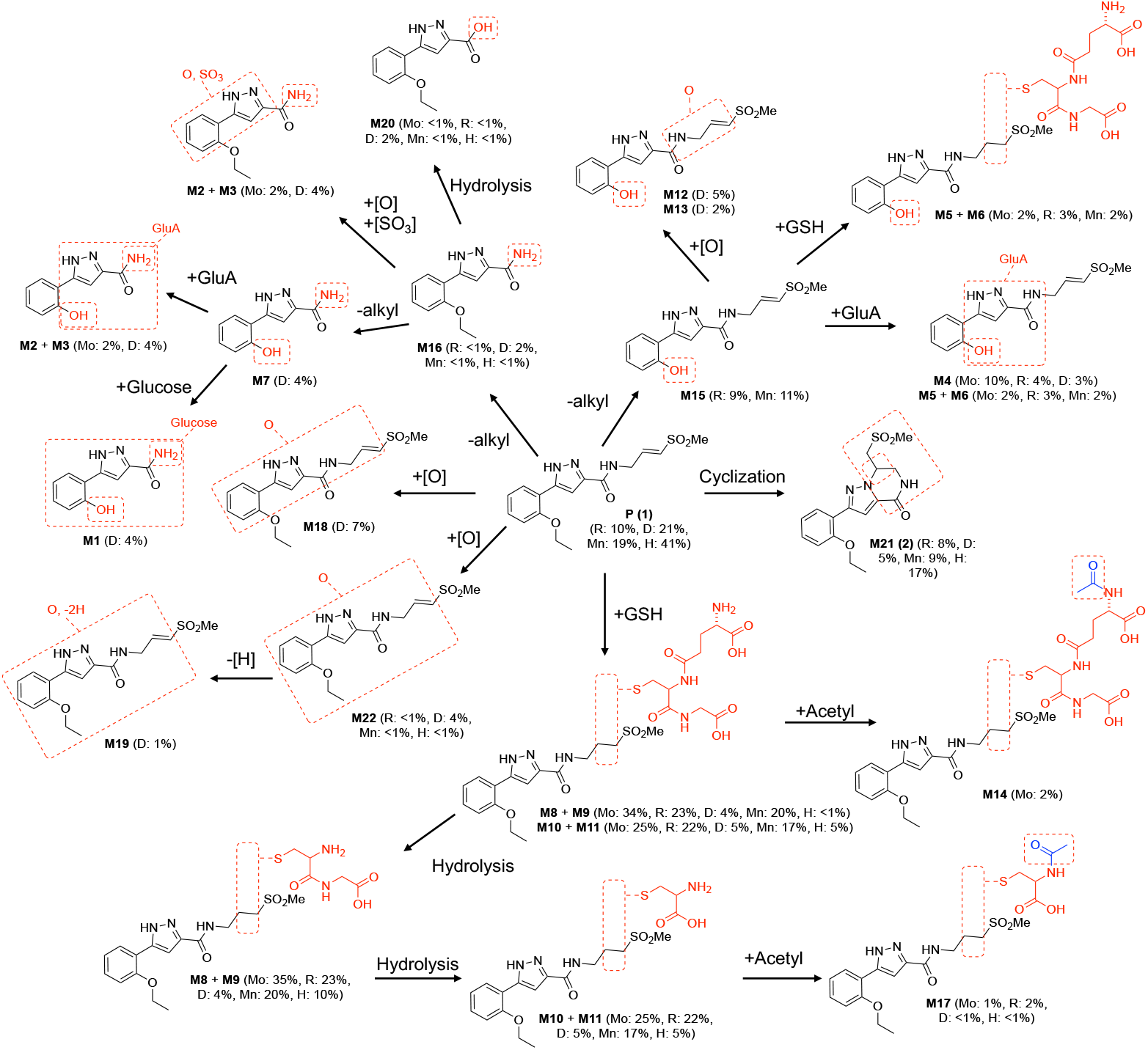
Metabolites of **1** in Primary Hepatocytes. Mo, mouse; R, rat; D, dog; Mn, monkey; H, human.

Since the rate of metabolism of **1** was dramatically different between mouse and human hepatocytes, we sought to determine whether species other than mice would be better suited for preclinical efficacy studies and allometric dose projections. Notably rat and monkey hepatocytes had intermediate rates of metabolism with >2:1 ratio of GSH conjugation to P450 oxidation (Table 4). In contrast, dog hepatocytes had higher levels of oxidative metabolism, which was only seen at very low levels in human hepatocytes. We proceeded to determine the pharmacokinetics of **1** in rats, dogs, and monkeys (Table 6 and File S2).

**Table 6.** Rat, Dog, and Monkey Pharmacokinetics of 1^a^.

|  | i.v. (10 mg/kg) |  |  |  | p.o. (30 mg/kg) |  |  |
| --- | --- | --- | --- | --- | --- | --- | --- |
|  | AUC<sub>last</sub><br>(h*ng/mL) | Vd<sub>ss</sub><br>(L/kg) | t<sub>1/2</sub><br>(h) | CL<sub>s</sub><br>(mL/min/kg) | C<sub>max</sub><br>(ng/mL) | AUC<sub>last</sub><br>(h*ng/mL) | F<br>(%) |
| Rat<sup>b</sup> | 3800 | 0.6 | 0.8 | 45 | 2900 | 4500 | 40 |
| Dog<sup>c</sup> | 6300 | 0.8 | 0.4 | 28 | 7100 | 9300 | 49 |
| Monkey<sup>d</sup> | 3000 | 1.9 | 0.6 | 56 | 2800 | 5600 | 63 |

<sup>a</sup>Based on plasma concentrations. All data n = 3. <sup>b</sup>Male SD rats. <sup>c</sup>Male beagle dogs. <sup>d</sup>Male cynomolgus monkeys.

Intravenous dosing showed that systemic clearance of **1** was in the moderate to high range in all three species and half-lives were <1 h (Table 6). Although systemic clearance from the plasma compartment was rapid, it was lower than in mice (Table 1), a result consistent with the cross-species primary hepatocyte data (Table 4). The results suggested GST-catalyzed GSH conjugation was still a major pathway of clearance in rats, dogs, and monkeys. Despite the high systemic clearance, the plasma profile following oral dosing at 30 mg/kg showed detectable levels of **1** up to 8 h post dosing in all three species (Figure 7). In SD rats, the oral bioavailability of **1** using AUClast was 40% with a rapid absorption phase and a slow elimination phase. In dogs, absorption was also rapid, but elimination was faster than in rats. The oral bioavailability in dogs was 49%. In cynomolgus monkeys, the oral bioavailability of **1** was 63% with a plasma profile that showed a slow absorption phase to a Cmax at 2h followed by a rapid elimination phase.

**Figure 7.**
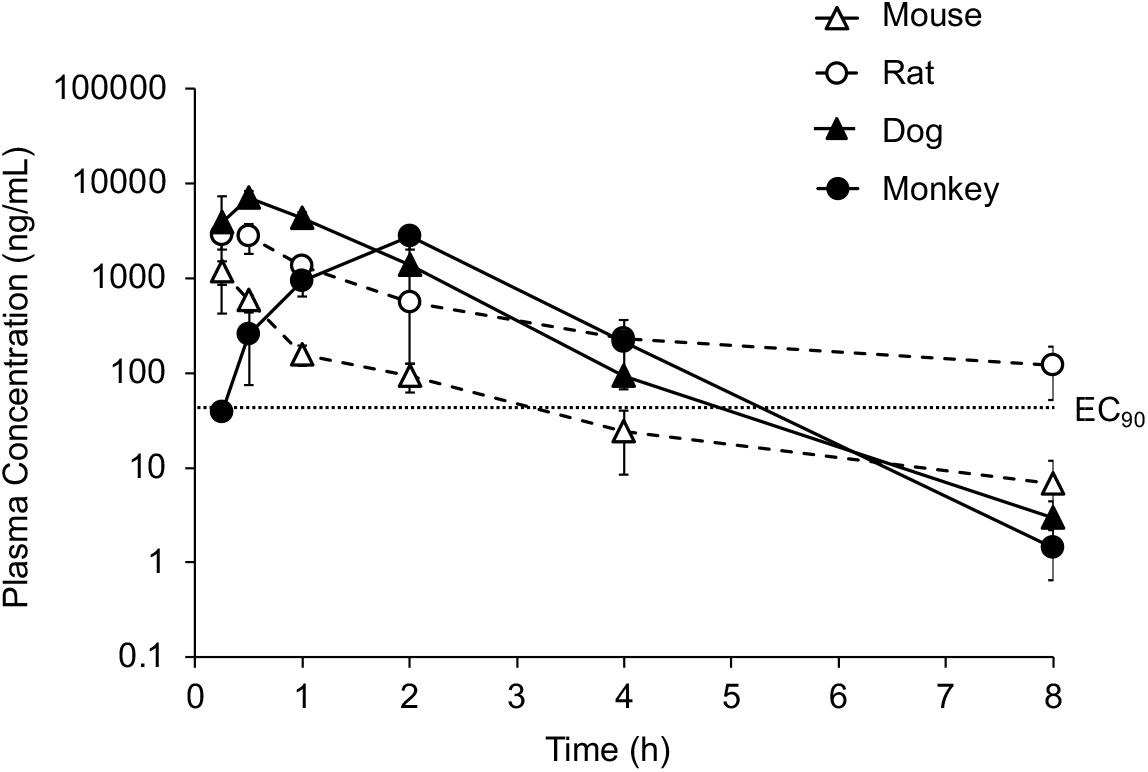
Cross-species Plasma Profile of **1**.Oral dosing of **1** at 30 mg/kg in four species from Tables 1 and 6. Dotted line indicates the EC90 for inhibition of CHIKV replication.

The pharmacokinetic profile of **1** in rats, dogs, and monkeys after a 30 mg/kg oral dose (Figure 7) revealed that plasma concentrations remaining above the CHIKV EC90 for >5 h compared to 3 h in mice.

### Human dose modeling

The cross-species data on intrinsic clearance from primary hepatocytes and systemic clearance data from *in vivo* pharmacokinetic studies in animals were used to begin to model the potential for oral dosing of **1** in human clinical studies (Table 7). Since data was not available to estimate the contribution of renal and other non-hepatic clearance, the calculations could only estimate the impact of hepatic clearance to drug concentrations. Thus, despite preclinical evidence that non-hepatic tissues may also contribute to drug clearance, our

**Table 7.** Predicted Hepatic Clearance of 1^a^.

|  | Mouse | Rat | Dog | Monkey | Human |
| --- | --- | --- | --- | --- | --- |
| $CL_{int, in vitro}$<br>($\mu\text{L}/\text{min}/10^6$ cells)<sup>b</sup> | 94–105 | 15 | 29 | 14 | 4.0–6.4 |
| Scaling factor<br>($10^6$ cells*$\text{mL}/\text{kg}*\mu\text{L}$) | 8.6 | 4.7 | 5.9 | 3.1 | 2.7 |
| Scaled $CL_{int, in vivo}$<br>($\text{mL}/\text{min}/\text{kg}$) | 805–900 | 52 | 170 | 31 | 9.8–16 |
| Calculated $CL_h$<br>($\text{L}/\text{h}$)<sup>d</sup> | 0.071–0.075 | 0.075 | 12.0 | 1.2 | 5.5–8.4 |
| Hepatic extraction | 0.59–0.62 | 0.069 | 0.48 | 0.10 | 0.063–0.097 |

<sup>a</sup>Equations listed in Table S2. <sup>b</sup>From Tables 2 and 4. <sup>c</sup>From Tables 1 and 6. HE: hepatic extraction ratio. <sup>d</sup> $CL_{int, in vivo}$ adjusted for $f_u$ (Table S1) and blood:plasma ratios (Table S3)

initial model assumed that systemic clearance was equal to hepatic clearance (CLs = CLh). Data from Phase I clinical studies to determine the half-life and volume of distribution of **1** will be required to build more sophisticated models. *In vitro*-*in vivo* extrapolation (IVIVE) scaling factors were calculated using the consensus values for hepatocellularity, liver weight, and liver blood flow collated by the human PK Prediction Working Group (Table S2).^12^ Despite recent concerns that cryopreserved hepatocytes may underestimate the rate of GST-catalyzed GSH conjugation,^13^ we found this not to be the case with metabolism of **1** (Table 4). IVIVE scaling using the data from primary hepatocytes was used to calculate CLint, *in vivo* (Table 7). Using a well-stirred model^14^ and assuming typical physiological parameters,^15^ the hepatic clearance (CLh) was calculated from the scaled CLint, *in vivo* across all five species. Since *in vivo* drug concentrations were measured in plasma, CLh was adjusted using blood to plasma ratio measured rats, monkeys and humans (Table S3) and the unbound drug using the fu value in mouse, rat, dog, and human blood (Table S1). The calculated CLh after adjusting for body weight ranged from ∼0.07 L/h in mice to 5.5–8.4 L/h in humans (Table 7). Comparison of CLh to hepatic blood flow^12^ yielded an estimate of hepatic extraction in each species, which was high for mouse, intermediate for dog, and low for rat, monkey and human.

In the mouse study, **1** demonstrated antiviral efficacy when the total plasma level was sustained above the EC90 (Figure 5). For human dose modeling, a target Css of 2 x EC90 (0.112 mg/L) was selected. This drug concentration would equate to a continuous i.v. infusion rate of 0.75–1.15 mg/h or a total daily i.v. dose of 18–28 mg based on the calculated CLh in humans (Table 7 and S2). To make a preliminary estimate of oral dosing regimens to use in human clinical trials the volume of distribution was set to 1.09 L/kg (76.8 L) using the mean of the values found in the four preclinical species. Assuming hepatic clearance as the only route of drug elimination, calculation of half-life as t1/2 = 0.093 x Vd / CLh resulted in a range of 6.3–9.8 h indicating that it would be possible to dose **1** b.i.d. in clinical studies. However, pharmacokinetic data from Phase I studies in humans, including the volume of distribution and half-life of **1**, will be needed to support calculation of the oral dose for antiviral efficacy. Furthermore, due to the irreversible inhibition of nsP2pro by **1**, it also remains to be determined whether the total peak^9^ or unbound trough^16^ drug concentration will be the determinant of clinical efficacy given that resynthesis of the target enzyme will depend on replication rate of the remaining virus.

## Conclusions

RA-0002034 (**1**), a potent covalent inhibitor of alphavirus nsP2 cysteine protease, demonstrated *in vivo* activity in a mouse model of CHIKV infection when dosed t.i.d. at 100 mg/kg. However, it is likely that the efficacy of **1** in mice was compromised by its high systemic clearance and short half-life. *In vitro* metabolism studies showed that **1** was a substrate for mouse GST, which was responsible for the extensive GSH conjugation of the vinyl sulfone warhead in hepatocytes and *in vivo*. Moreover, with i.v. dosing, total systemic clearance exceeded hepatic blood flow indicating that GST in both liver and peripheral tissues may contribute to the metabolism of **1** in mice.

Additional *in vitro* metabolism and *in vivo* pharmacokinetic studies showed that the clearance of **1** was species dependent. Surprisingly, systemic clearance in rats was only 1/4 of the rate seen in mice. Likewise in dogs and monkeys, *in vivo* clearance was lower than in mice. Studies using primary hepatocytes also demonstrated cross-species differences in the major pathway of metabolism. In mouse, rat, monkey, and human hepatocytes GSH conjugation was dominant over P450 oxidation, whereas in dog hepatocytes the situation was reversed with oxidation seen as the principal metabolic pathway. These insights into the species dependent metabolism of **1** will be invaluable for the design of new analogs with reduced systemic clearance that may have improved pharmacokinetic profiles.

Studies in primary hepatocytes and IVIVE predicted low hepatic extraction of **1** in rats, monkeys and humans. Using a model in which hepatic clearance was the only path for elimination of the drug, it was projected that a clinical b.i.d. dosing of **1** could be used to maintain its concentration at an appropriate level for antiviral activity. This calculation supported the continued progression of **1** toward clinical development as a direct acting anti-alphaviral drug to determine key pharmacokinetic parameters in Phase I studies in order to obtain an estimation of the effective antiviral dose.

## Experimental Section

### Pharmacokinetics

Male CD-1 mice (n = 3, 6−8 weeks, 20−30 g) were dosed by p.o. administration of **1** (30 mg/kg) in NMP/Solutol/PEG-400/normal saline (v/v/v/v, 10:5:30:55). Blood (0.03 mL) was collected from the dorsal metatarsal vein at 0.25, 0.5, 1, and 3 h time points. Blood from each sample was transferred into plastic microcentrifuge tubes containing K2EDTA, mixed well, and then placed in a cold box prior to centrifugation. Blood samples were centrifuged at 4000g for 5 min at 4 °C to obtain plasma and then stored at −75 °C prior to analysis. Concentrations of **1** in the plasma samples were determined using an AB API 5500+ LC-MS/MS instrument fitted with a HALO 160A C18 column (2.7 μm, 2.1 × 50 mm) using a mobile phase of 5−95% MeCN in H2O with 0.1% formic acid. Pharmacokinetic parameters were calculated from the mean plasma concentration versus time by a non-compartmental model using WinNonlin 8.3 (Phoenix) to determine Cmax, AUClast, t1/2, CLint.

For co-dosing studies, male C57BL/6 mice (n = 3, 6−8 weeks, 20−30 g) were dosed by p.o. administration of **1** (30 mg/kg p.o.) as a 10 mL/kg volume of a 3 mg/mL solution in NMP/Solutol/PEG-400/normal saline (v/v/v/v, 10:5:30:55). EA (10 or 30 mg/kg p.o.) was administered as a 5 mL/kg volume of a 6 mg/mL solution in NMP/PEG-400/water (v/v/v, 10:60:30) as a 6 h pretreatment and again at the time of dosing of **1**. 1-ABT (100 mg/kg p.o.) was administered as 10 mL/kg volume of a 10 mg/mL solution in saline as a 2 h pretreatment. Blood (0.03 mL) was collected from the dorsal metatarsal vein at 0.5, 1, 3, and 5 h time points and the level of **1** in plasma was determined by LC-MS/MS analysis.

For t.i.d. studies, male C57BL/6 mice (n = 3, 6−8 weeks, 20−30 g) were dosed by p.o. administration of **1** (100 mg/kg) in NMP/Solutol/PEG-400/normal saline (v/v/v/v, 10:5:30:55) at 0, 8, and 16 h respectively. Blood (0.03 mL) was collected from the dorsal metatarsal vein at 0.5, 1, 2, 4, 8, 10, 16, 18, and 24 h time points post first doing and the level of **1** in plasma was determined by LC-MS/MS analyses.

Male Sprague-Dawley rats (n = 3, 6–8 weeks, 200–300 g) were dosed with **1** by i.v administration (10 mg/kg, 2 mg/mL) or p.o. administration (30 mg/kg, 6 mg/mL) in NMP/Solutol/PEG-400/normal saline (v/v/v/v, 10:5:30:55). Male beagle dogs (n = 3, 0.7–3 y, 6– 13 kg) were dosed with **1** by i.v administration (10 mg/kg, 10 mg/mL) or p.o. administration (30 mg/kg, 6 mg/mL) in PEG400/TPGS/20%HP-bet-CD in water+1N NaOH to make pH 3–4 (v/v/v/v, 10:5:85). Male cynomolgus monkeys (n = 3, 2–5 y, 2–6 kg) were dosed with **1** by i.v administration (10 mg/kg, 5 mg/mL) or p.o. administration (30 mg/kg, 6 mg/mL) in DMSO/PEG-400/Water (v/v/v, 5:40:55). Serial blood samples were collected by jugular vein cannula (rats) or venipuncture of peripheral veins (dogs and monkeys) at 0.5, 1, 2, 4, 8, 10, 16, 18, and 24 h after the first dose and the plasma level of **1** was determined by LC-MS/MS analyses.

### Mouse whole blood stability

A 1 mM working solution of **1** was prepared in DMSO, 1 mM working solution of control compound propantheline was prepared in MeCN. 4 µL of working solution was spiked to 796 µL of pre-incubated whole blood from CD-1 mice to reach a final concentration of 5 µM. The final concentration of solvent was 0.5%. 50 µL aliquots of the spiked whole blood were added into new tubes for 15, 30, 60, and 120 min timepoints and incubated at 37 °C water bath with shaking at 60 rpm. The assay was performed in duplicate. The reaction was stopped by adding 400 µL of cold MeCN containing IS (100 nM Alprazolam, 500 nM Labetalol and 2 µM Ketoprofen). Samples were centrifuged at 3,220 g for 30 min, and 100 μL of the supernatant mixed with 100 μL of ultrapure water for LC-MS/MS analysis. Peak areas were determined from extracted ion chromatograms, and the slope value, *k*, was determined by linear regression of the natural logarithm of the remaining percentage of the parent drug vs. incubation time curve. The *in vitro* t1/2 was determined from the slope value using the relationship: *in vitro* t1/2 = –0.693/k, where k is the rate constant.

### Liver microsome stability

A 10 mM DMSO stock of **1** was diluted to 0.5 mM with DMSO and again to 0.1 mM with MeCN. The resulting solution contained 0.1 mM of **1** in 20% DMSO/80% MeCN. Male CD-1 mouse liver microsomes (Xenotech, lot no. 2410002), supplied as 20 mg/mL protein concentration were used for the assay. A reaction plate was prepared by adding 691.25 μL, pre-warmed (37 °C) microsomal solution (0.63 mg/mL protein in 100 mM KPO4 with mM EDTA) to a 96-well plate and maintained at 37 °C. The diluted 0.1 mM **1** (8.75 μL) was added to the microsomal solution in the reaction plate and mixed thoroughly by repeated pipetting. The resulting solutions were pre-incubated for 5 min at 37 °C. For the t = 0, 5, 15, 30, 60 min incubation plates, NADPH (95 μL) was added to the each well, the plate was sealed and incubated at 37 °C for the incubation period. An aliquot (100 μL) was removed from each well at the desired time point and dispensed into a well of a 96-well plate. The reaction was quenched using 200 μL of cold (4 °C) MeOH. All plates were sealed, vortexed, and centrifuged at 3000 rpm, 4 °C for 15 min, and the supernatants were transferred for analysis by LC-TOFMS. The supernatant was injected onto an AQUASIL C18 column and eluted using a fast-generic gradient program. TOFMS data was acquired using Agilent 6538 Ultra High Accuracy TOFMS in extended dynamic range (*m/z* 100–1000) using generic MS conditions in positive mode. Following data acquisition, exact mass extraction and peak integration were performed using MassHunter Software (Agilent Technologies). The stability of **1** was calculated as the percent remaining of the unchanged parent compound at each time point relative to the peak area at t = 0 min. Peak areas were determined from extracted ion chromatograms, and the slope value, *k*, was determined by linear regression of the natural logarithm of the remaining percentage of the parent drug vs incubation time curve. The t1/2 was determined from the slope value using the relationship: t1/2 = –0.693/*k*, where *k* is the rate constant. The intrinsic clearance (CLint in μL/min/mg) was calculated using the relationship CLint = kV/N where V is the incubation volume and N is the amount of protein per well.

### Hepatocyte stability

Cryopreserved hepatocytes from mouse (TPCS, cat. no. CMH-100CD-SQ, lot no. CMH100CD-V01422 and CMH100CD-V01699, pooled male CD-1), rat (BioIVT, cat. no. M00005, lot no. FYT, pooled male Sprague-Dawley), dog (BioIVT, cat. no. M00205, lot no. BQU, pooled male beagle), monkey (RILD, cat. no. HP-SXH-02M, lot no. GRLG, pooled male cynomolgus), and human (BioIVT, cat. no. X008001, lots no. QZW and YIT, 10 pooled donors each) were used. 10 mM stock solutions of **1** and positive control verapamil were prepared in DMSO. Thawing medium and supplement incubation medium (serum-free) were placed in a 37 °C water bath for at least 15 min prior to use. Stock solutions were diluted to 100 μM by combining 198 μL of 1:1 MeCN/water and 2 μL of 10 mM stock solution. Vials of cryopreserved hepatocytes were removed from cryogenic storage and thawed in a 37 °C water bath with gentle shaking until all ice crystals had dissolved. The contents were poured into the 50 mL thawing medium conical tube and centrifuged at 100 g for 10 min at room temperature. Thawing medium was aspirated, and hepatocytes were re-suspended with serum-free incubation medium to yield ∼1.5 × 10^6^ cells/mL. Cell viability and density were counted using AO/PI fluorescence staining and then diluted with serum-free incubation medium to a working cell density of 0.5×10^6^ viable cells/mL. A portion of the hepatocytes at 0.5×10^6^ viable cells/mL was boiled for 5 min prior to adding to the plate as negative control. Aliquots of 198 μL hepatocytes were dispensed into each well of a 96-well non-coated plate. For the inhibitor plate, 0.5 μL of 80 mM EA (0.2 mM) or 400 mM 1-ABT (1 mM) were added. The plate was placed in the incubator for approximately 20 min. Aliquots of 2 μL of 100 μM **1** (1 μM) and positive control were added into respective wells of the non-coated 96-well plate to start the reaction. The assay was performed in duplicate. The plate was incubated for the designed time points. 25 μL of contents were transferred and mixed with 6 volumes (150 μL) of cold MeCN with IS (100 nM alprazolam, 200 nM labetalol, 200 nM caffeine and 200 nM diclofenac) to terminate the reaction at time points of 0, 15, 30, 60, 90 and 120 min. Samples were centrifuged for 45 min at 3,220 g and an aliquot of 100 µL of the supernatant was diluted by 100 µL ultra-pure H2O, and the mixture was used for LC-MS/MS analysis. Peak areas of **1** were determined from extracted ion chromatograms. The *in vitro* t1/2 was determined from the slope value using the relationship: *in vitro* t1/2 = –0.693/k, where k is the rate constant. The intrinsic clearance in μL/min/10^6^ cells was calculated using the relationship CLint = kV/N where V is the incubation volume (0.2 mL) and N is the number of hepatocytes per well (0.1 × 10^6^ cells).

### GSH capture

The GSH reactivity of **1** was determined using a modification of the reported protocol.^7^ Recombinant mouse GST Mu 1 protein (mGSTM1) was purchased from AssayPro (catalog no. PMG30413, size: 20 ug, >95% purity by SDS-PAGE). Recombinant human GST A1 protein (hGSTA1) was purchased from Abcam (catalog no. ab276644, >90% purity by SDS-PAGE). A 100 mM solution of GSH (Sigma Aldrich cat. no. G4251) was prepared in a pH 7.4 phosphate buffer. A 10 mM DMSO solution of compound **1** was diluted in phosphate buffer to give a solution at 1000 µM. Three independent experiments were performed: (i) with **1** alone, (ii) with **1** and mGSTM1; (iii) with **1** and hGSTA1. Experiment (i): at time zero (t = 0), 5 µL of the 1000 µM solution of **1** was added to an Eppendorf tube containing 90 µL phosphate buffer and 5 µL of 100 mM GSH solution. Experiment (ii): at time zero (t = 0), 5 µL of the 1000 µM solution of **1** and 1 µL of mGSTM1 (1 mg/mL) were added to an Eppendorf tube containing 89 µL phosphate buffer and 5 µL of 100 mM GSH solution. Experiment (iii): at time zero (t = 0), 5 µL of the 1000 µM solution of **1** and 1 µL of hGSTA1 (1 mg/mL) were added to an Eppendorf tube containing 89 µL phosphate buffer and 5 µL of 100 mM GSH solution. The final concentrations of **1**, GSH, and mGSTM1/hGSTA1 were maintained at 50 µM, 5 mM, and 10,000 ng/mL respectively. The Eppendorf tubes were vortexed, and then the samples were transferred to high recovery autosampler vials for LCMS analysis. Analysis was performed at 0, 2, 4, 6, 8, 10, 12 h time points, and the percentages of GSH adduct formation were calculated using Agilent LCMS software (OpenLab CDS Version 2.7).

### Plasma protein binding

Frozen plasma that had been stored at -80°C was thawed in a 37°C water bath. Working solutions of test compound **1** and the control ketoconazole were prepared in DMSO at 200 μM and then spiked into plasma. The final concentration of the compounds was 1 μM with 0.5% DMSO. The dialysis membranes were soaked in ultrapure water for 60 min to separate strips, in 20% ethanol for 20 min, and finally in dialysis buffer for 20 min. Each dialysis cell was filled with 150 μL of plasma sample and dialyzed against equal volume of PBS buffer. The assay was performed in duplicate. The dialysis plate was sealed and incubated at 37°C with 5% CO2 at 100 rpm for 6 h. 50 μL of samples from both buffer and plasma chambers were transferred to a 96-well plate. 50 μL of plasma was added to each buffer sample and supplemented with an equal volume of PBS. A 400 μL of quench solution (200 nM labetalol, 100 nM tolbutamide, and 100 nM ketoprofen in acetonitrile) was added to precipitate protein and release compounds. Samples were vortexed for 2 min and centrifuged for 30 min at 3,220 g. 100 µL of the supernatant was diluted with 100 µL of ultra-pure H2O, and the mixture was used for LC-MS/MS analysis.

### Blood to plasma Ratio

Blood samples were collected freshly before the experiments. Stock solutions of test compound **1** in DMSO and control chloroquine phosphate in ultrapure water at 10 mM were prepared and then diluted 50x with DMSO to working concentration of 0.2 mM. Plasma was obtained from whole blood by centrifugation at 10,000 g (4°C) for 10 min. 2 μL of the compound solution was added to 398 μL plasma and blood to achieve a final concentration of 1 μM. The assay was performed in duplicate. The sample tubes were incubated at 37°C with shaking for 60 min. The blood samples were spun for 10 min at 10,000 g (37°C) and the plasma samples stored at 37°C. 100 μL aliquots of plasma from the centrifuged whole blood samples and reference plasma samples were placed into tubes. 400 μL of quench solution (200 nM labetalol, 100 nM tolbutamide, and 100 nM ketoprofen in acetonitrile) was added to precipitate protein. The mixture was vortexed for 5 minutes and centrifuged for 15 min at 20,000 g. 100 μL of the supernatant was transferred to a new plate and diluted with 100 μL or 200 μL water according to the LC/MS signal response and peak shape. Samples were mixed well and analyzed using LC-MS/MS.

### Metabolite identification

**1** was administered p.o. at 30 mg/kg dose to male CD-1 mice (n = 3). Plasma samples were collected at 0.25, 0.5, 1, 3, and 5 h intervals post dose and pooled. For each time point, a calculated volume of 10 µL was taken from individual samples. These samples were pooled based on an equal volume pooling scheme. Specifically, for each time point, 10 µL was pooled from three mice, resulting in a total pooled volume of 30 µL per time point. The cumulative pooled volume across all five time points was 150 µL (50 µL x 3). 150 μL of pooled volume was mixed with 600 μL of MeCN. The samples were vortex-mixed for 100 seconds before centrifugation at 16,000 g for 30 min at 4 °C to precipitate proteins. 700 μL of the supernatants were transferred to clean tubes and placed in the evaporator under steady stream of nitrogen at room temperature until dry. The dried residues were reconstituted with 100 μL of 20:80 (v/v) MeCN/water and vortexed for proper mixing. The mixtures were centrifuged for 15 min at 16,000 g to precipitate the protein and injected into an LC-UV-MS/MS. Samples for metabolite identification and profiling were analyzed using a Vanquish UHPLC system, equipped with Diode

Array Detector HL and Orbitrap Exploris 120 (Thermo Fisher Scientific, USA). A XSelect HSS T3, 100 × 2.1 mm, 2.5 µm HPLC column (Waters) was used for separation of analytes. The mobile phase was a gradient of water, containing 0.1% formic acid (A) and MeCN, containing 0.1% formic acid (B). The flow rate was 500 µL/min. The mobile phase composition began at 5% B held for 0– 1.5 min, 5–50% B at 1.5–9 min, 50–100% B at 9–12 min, 100% B at 12–14 min, 100–5% B at 14–14.3 min, and finally 14.3–15 min re-equilibration period at 5% B. UV absorbance was monitored with a photodiode array detector from 220–400 nm. The mass spectrometer was operated in both negative and positive-ion modes with an electrospray ion source potential of 3.5 kV, and source heater temperature of 350 °C. Curtain, GS1 and GS2 gas flow settings were 25, 30, and 35 units, respectively. Full scan mass spectra were acquired over the range 120–1200 *m/z* at a constant resolving power of ∼ 60,000. Metabolites were characterized by comparison of mass spectral fragmentation patterns with that for compound 1 (see Figure 3 and Figure S1–S9 for structural assignments of metabolites).

For primary hepatocytes studies, **1** (10 µM) was administered to mouse (male CD-1, TPCS cat. no. CMH-100CD-SQ, lot no. CMH100CD-V01599), rat (male Sprague-Dawley, TPCS cat. no. CRH-100SD-SQ, lot no. CRH100SD-V01199); dog (male Beagle, BioIVT cat. no. M00205, lot no. BQU), monkey (male cynomolgus, RILD cat. no. HP-SXH-02M, lot no. SLFS), and human (mixed, liverpool^®^20-donor, BioIVT cat. no. X008000, lot no. ASW) hepatocytes, at a cell density of 1.0 × 10^6^ cells/mL. The medium used for the experiment was WEM (Williams’ Medium E) supplemented with GlutaMAX. Samples were incubated at 37 °C for three time points: 0, 1, and 2 h. The total volume for each sample was 200 µL. Incubations were quenched with 3 volumes of MeCN (0.1% formic acid) followed by centrifugation for 30 min at 16,000 g, and then 400 µL supernatants were transferred to clean tubes and place in the evaporator under steady stream of nitrogen at room temperature until dry. The dried residues were reconstituted with 100 µL of 1:4 (v/v) MeCN/H2O solution and vortexed for proper mixing and injected into LC-UV-MS/MS. for metabolite identification.

### Mouse antiviral study

Sets of 5 six week old male C57Bl/6J mice were treated p.o. with 100 mg/kg **1** in 100 μL diluent (10% NMP, 5% solutol, 30% PEG400, 55% saline) or diluent alone per every 8 h, with treatment initiated at 8 h prior to infection with 10^3^ PFU of the SL15649 strain of CHIKV virus through the subcutaneous route in the left rear footpad.^17^ Mice were euthanized at 48 h post infection by isoflurane overdose followed by cardiac puncture, and viral loads in the serum, spleen, right and left quadriceps, and right and left rear ankle/foot by plaque assay using established protocols.^18^

## Supporting Information

Table S1: Plasma Protein Binding of **1**; Table S2: Equations for Allometric Scaling; Table S3: Blood to Plasma Ratios of **1**; Figure S1: MS fragmentation of metabolites M1–18 from mouse plasma; Figure S2: MS fragmentation of metabolites M1–22 from hepatocytes; File S1: Pharmacokinetic data – mouse; File S2: Pharmacokinetic data – rat, dog, monkey,

## Acknowledgements

The authors gratefully acknowledge the support of Pharmaron for conducting *in vitro* metabolism and *in vivo* pharmacokinetic studies. Lillian Chang, Kara Carter, and Jeremy Travins from the READDI-AViDD Center scientific advisory board provided valuable insights and guidance for the preclinical studies of **1**.

