## Supporting Information for "Species Dependent Metabolism of a Covalent nsP2 Protease Inhibitor with *in Vivo* Anti-alphaviral Activity"

Table of Contents


| | Description | Page |
|---|---|---|
| Table S1 | Plasma Protein Binding of **1** | S2 |
| Table S2 | Equations for Allometric Scaling | S3 |
| Table S3 | Blood to Plasma Ratios of **1** | S4 |
| Figure S1 | MS fragmentation of metabolites M1–18 from mouse plasma | S5–13 |
| Figure S2 | MS fragmentation of metabolites M1–22 from hepatocytes | S14–21 |
| File S1 | Pharmacokinetic data – mouse | |
| File S2 | Pharmacokinetic data – rat, dog, monkey | |

**Table S1. Plasma Protein Binding of 1[a]**

| Species | Protein binding (%) | $f_u$ |
| --- | --- | --- |
| Mouse | 82 | 0.18 |
| Rat | 89 | 0.11 |
| Dog | 80 | 0.20 |
| Human | 86 | 0.14 |

[a] Plasma protein binding was determined by equilibrium dialysis. For monkey, the human value used.

**Table S2. Equations for Allometric Scaling**

| Calculated Value | Equation |
|---|---|
| Systemic Clearance | $CL_s = \frac{F \times X_0}{AUC_{PO}}, where\ F = \frac{AUC_{PO}}{AUC_{IV}} \times \frac{X_{0,IV}}{X_{0,PO}}$ |
| Intrinsic Clearance[a] | $CL_{int,in\ vivo} = \frac{hepatocellularity\ (\frac{10^6 cells}{g\ liver}) \times \frac{liver\ weight\ (g)}{kg\ body\ weight\ (kg)} \times CL_{int,in\ vitro}(\mu L/min/10^6 cells)}{fu_{in\ vitro}}$ |
| IVIVE Scaling Factor[b] | $Scaling\ factor = hepatocellularity\ (10^6 cells/g\ liver) \times liver\ weight\ (g/kg) \times \frac{1\ mL}{1000\ uL}$ |
| Blood Clearance | $CL_{int,blood} = CL_{int,plasma} \times \frac{C_{plasma}}{C_{blood}}$ |
| Hepatic Clearance | $CL_h = \frac{Q_{blood} \times fu \times CL_{int,in\ vivo}}{Q_{blood} + fu \times CL_{int,in\ vivo}}$ |
| Hepatic Extraction | $E_{hepatic} = \frac{\frac{Q_{blood} \times fu \times CL_{int,in\ vivo}}{Q_{blood} + fu \times CL_{int,in\ vivo}}}{Q_{blood}} = \frac{CL_h}{Q_{blood}}$ |
| i.v. Infusion Rate | $k_0 = infusion\ rate = C_{ss} \times CL$ |

[a]Calculations assumed $fu_{in\ vitro} = 1$

[b]Values from Petersson, C., Zhou, X., Berghausen, J., Cebrian, D., Davies, M., DeMent, K., & Baker, J. (2022). Current approaches for predicting human PK for small molecule development candidates: findings from the IQ human PK prediction working group survey. The AAPS journal, 24(5), 85. https://doi.org/10.1208/s12248-022-00735-9

**Table S3. Blood to Plasma Ratios of 1**

| | Blood:plasma ratio ($K_{b/p}$)[a] |
|---|---|
| Rat | 1.37 |
| Monkey | 1.41 |
| Human | 1.12 |

[a]For Mouse and Dog a nominal value of 1.0 was used in IVIVE calculations based on measured values in the other species.

**Figure S1.** MS fragmentation of metabolites M1–18 from mouse plasma

**M1**

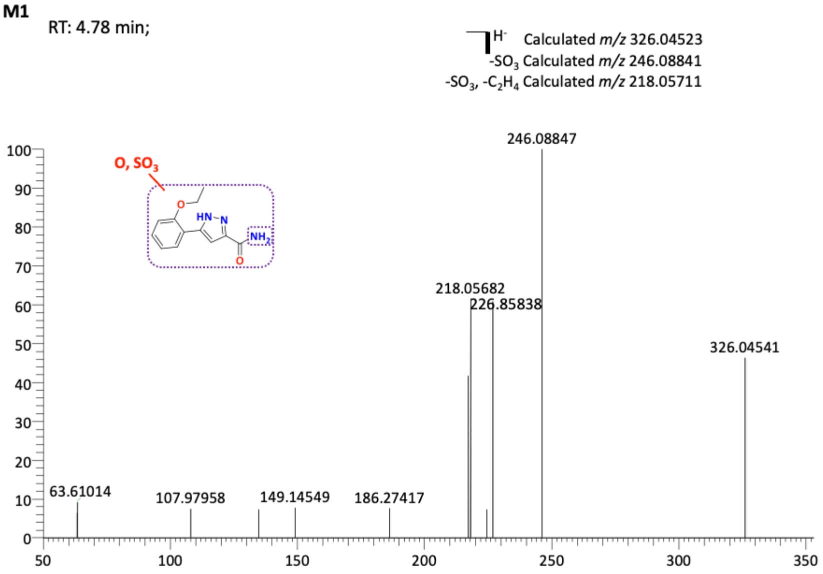


**M2**

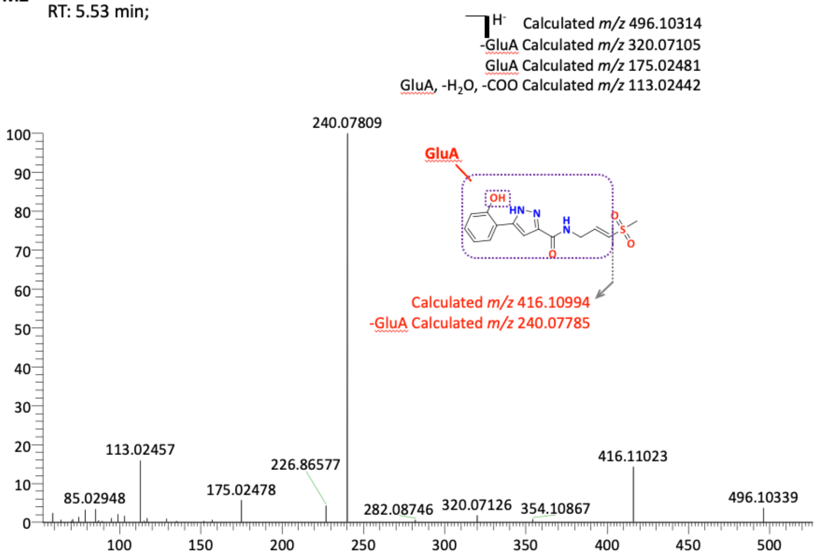

**M3**

RT: 5.60 min;

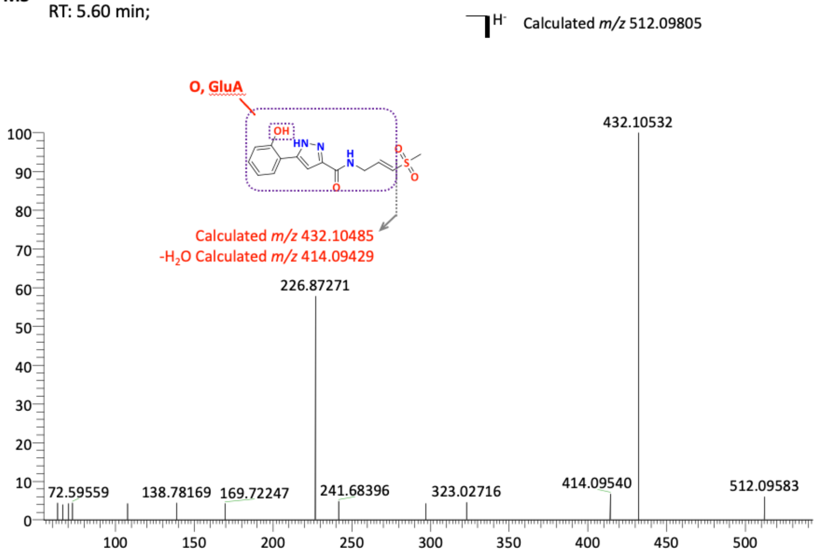


**M4**

RT: 5.62 min;

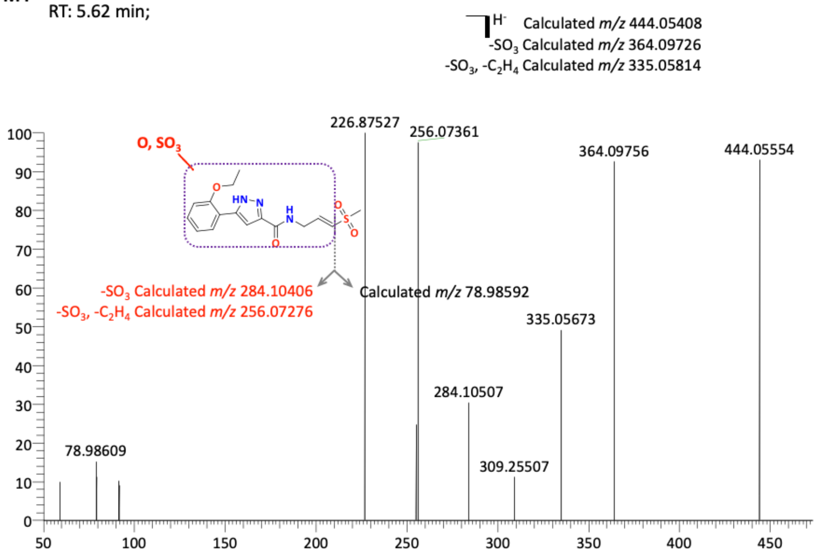

**M5**

RT: 5.72 min;

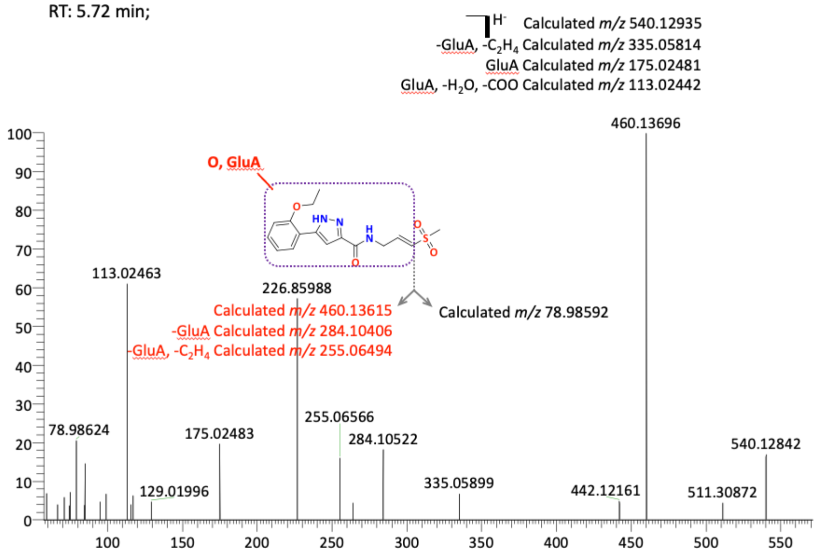


**M6**

RT: 6.06 min;

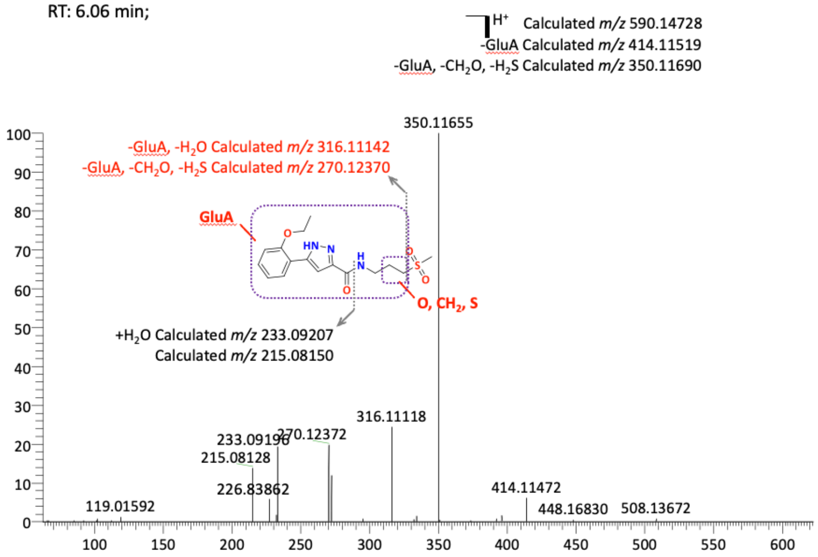

**M7**

RT: 6.16 min;

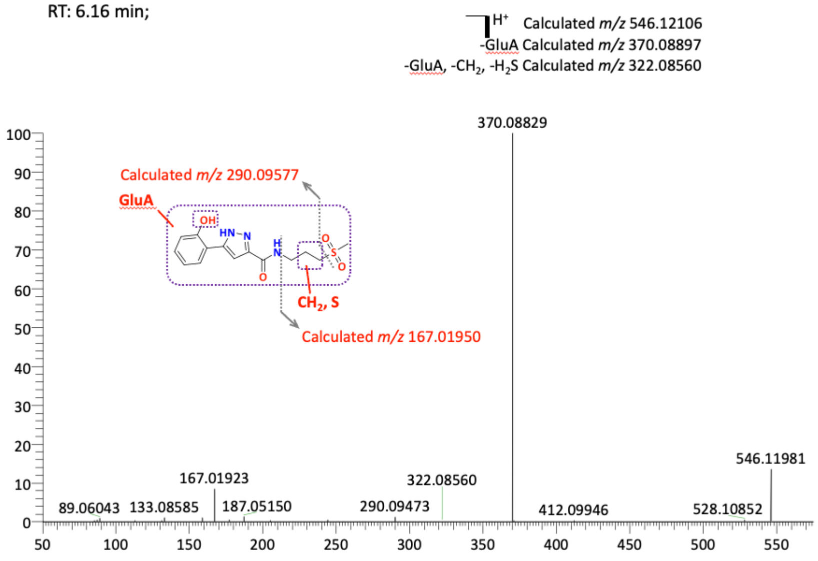


**M8**

RT: 6.29 min;

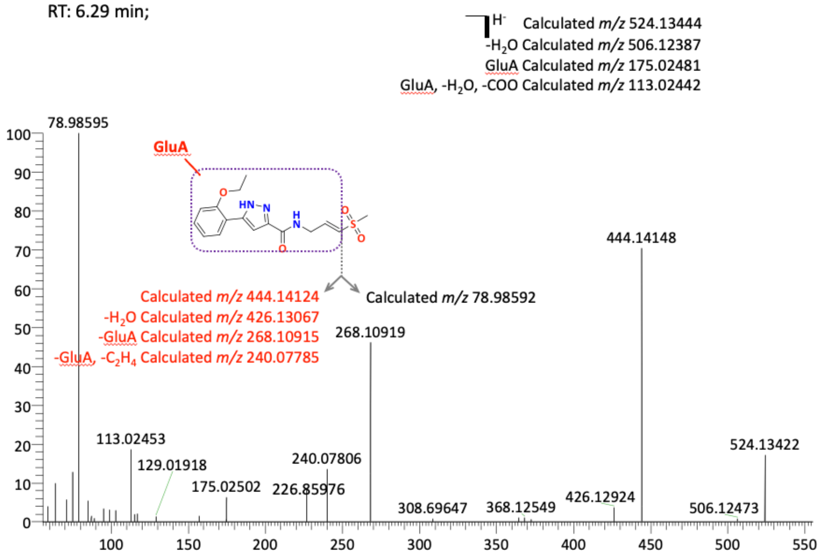

**M9**

RT: 6.50 min;

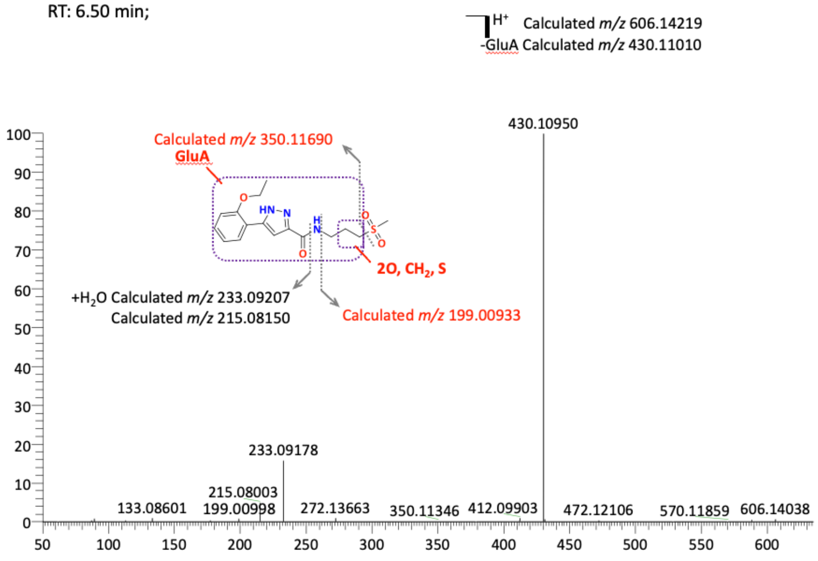


**M10**

RT: 6.70 min;

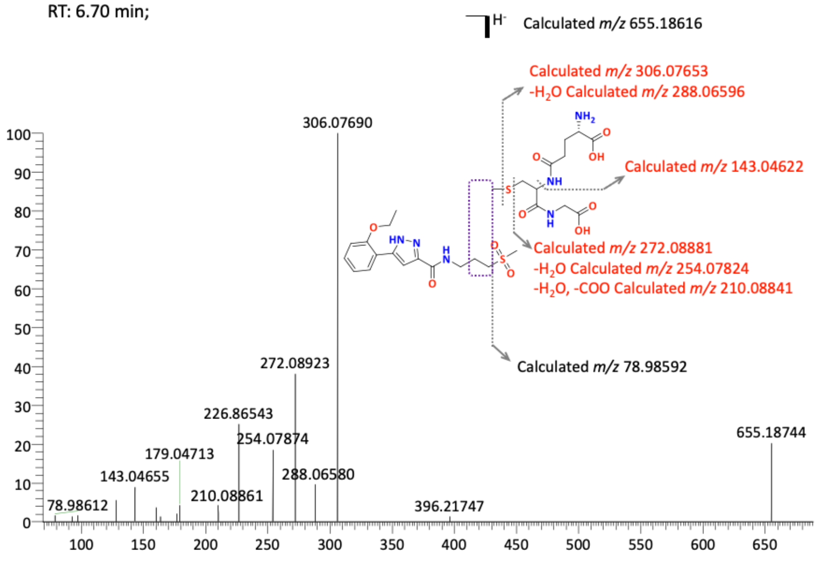

**M11**

RT: 6.74 min;

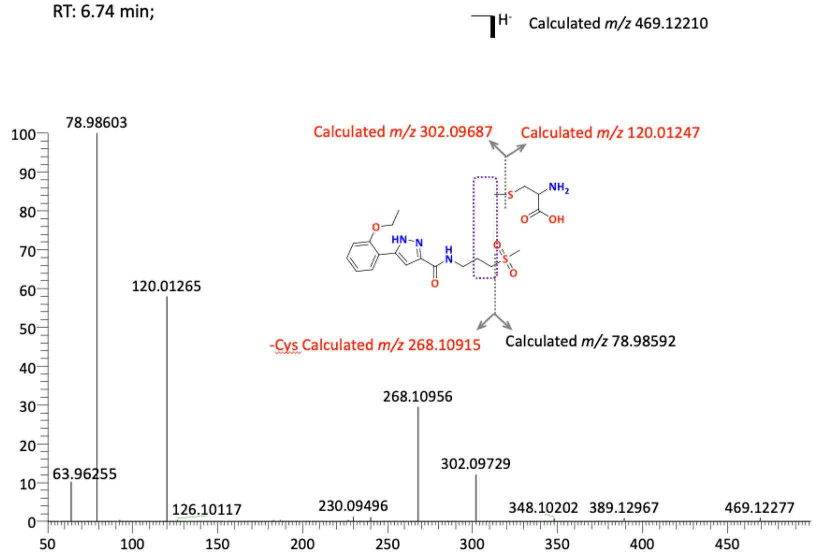


**M12**

RT: 6.81 min;

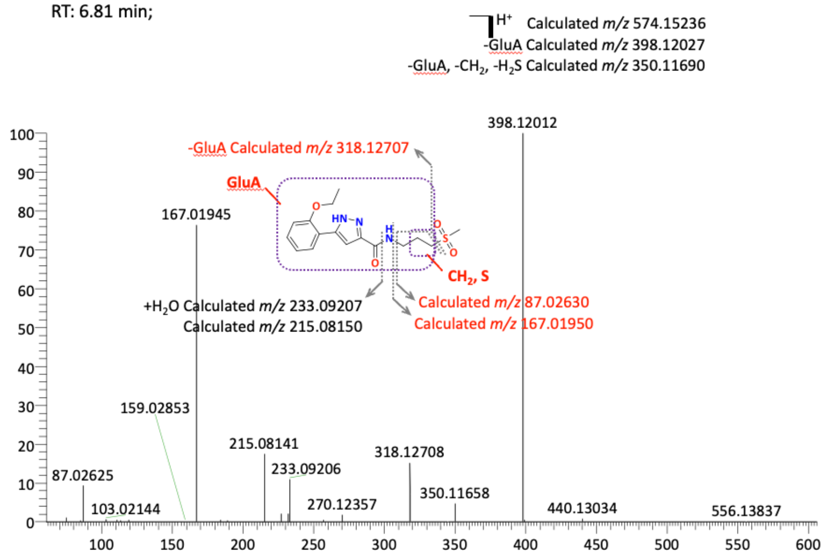

**M13**

RT: 7.02 min;

H⁻ Calculated *m/z* 460.13615

-GluA Calculated *m/z* 284.10406

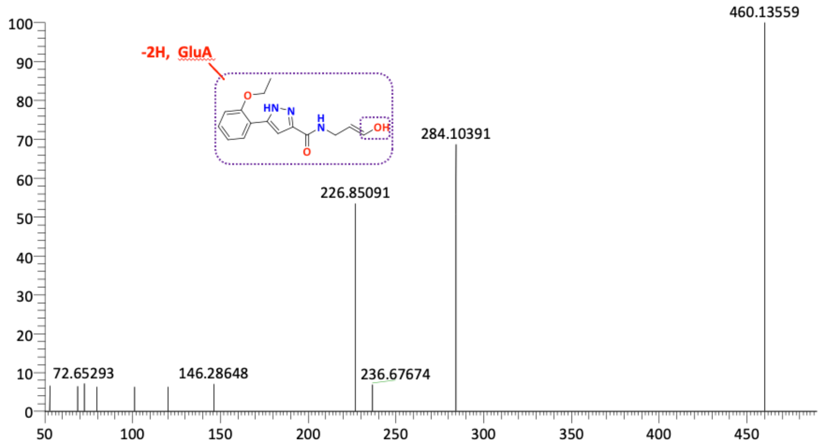


**M14**

RT: 7.49 min;

$H^+$ Calculated *m/z* 232.10805

$-NH_3$, $+H_2O$ Calculated *m/z* 233.09207

$-NH_3$ Calculated *m/z* 215.08150

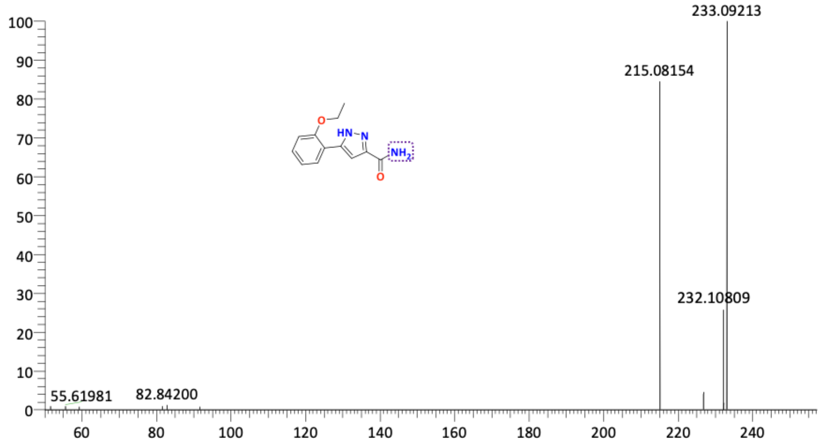

**M15**

RT: 7.75 min;

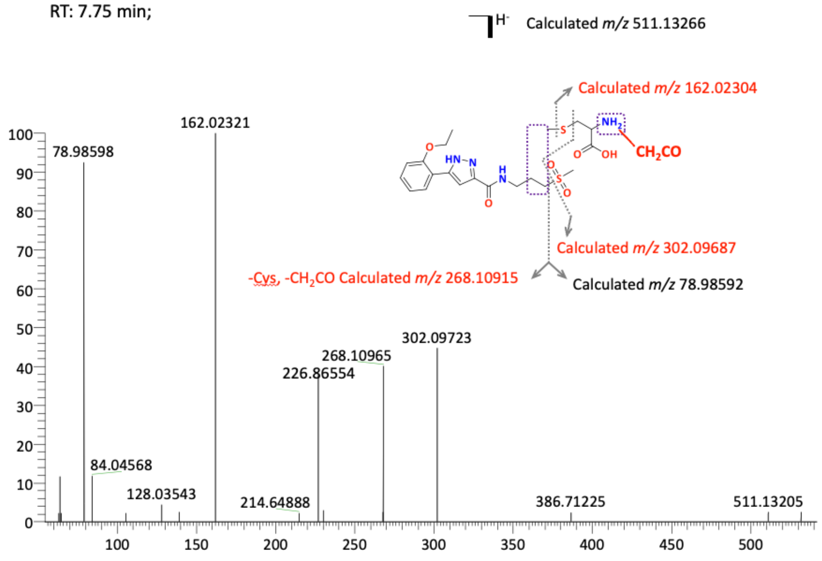


**M16**

RT: 8.06 min;

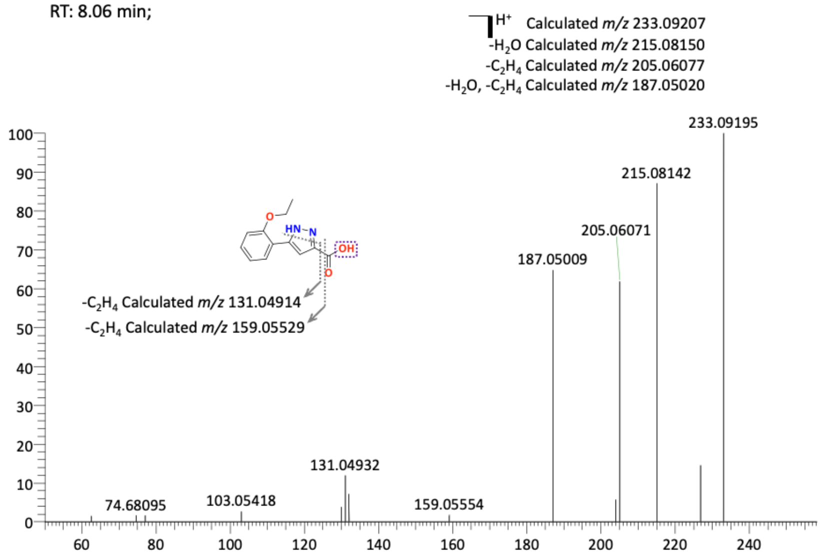

**M17**

RT: 8.34 min;

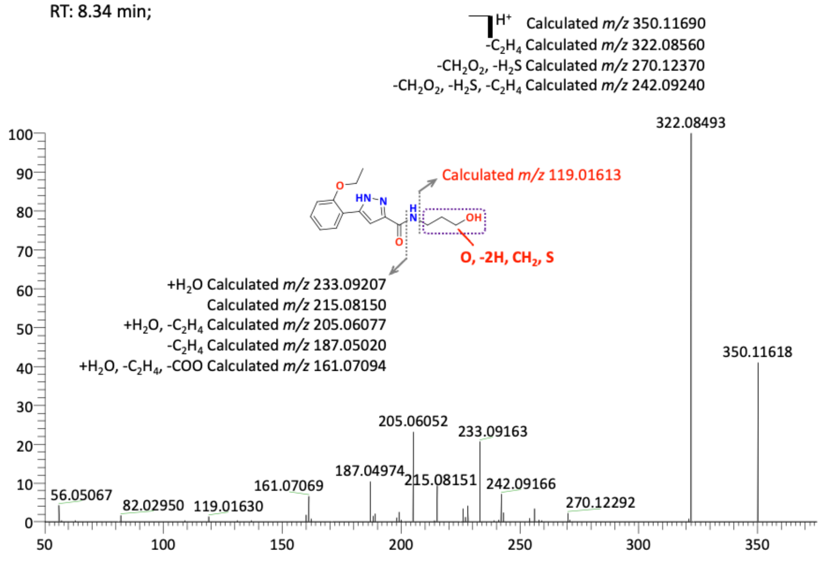


**M18**

RT: 8.97 min;

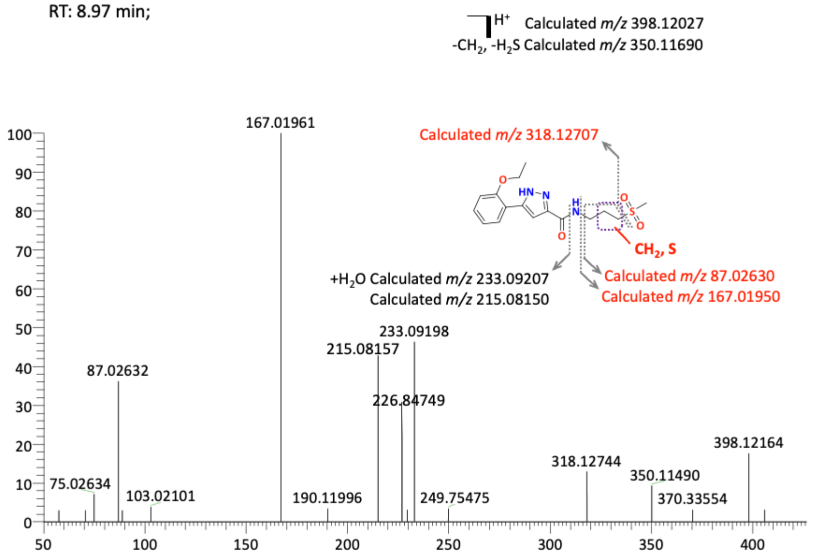

**Figure S2.** MS fragmentation of metabolites M1–22 from hepatocytes

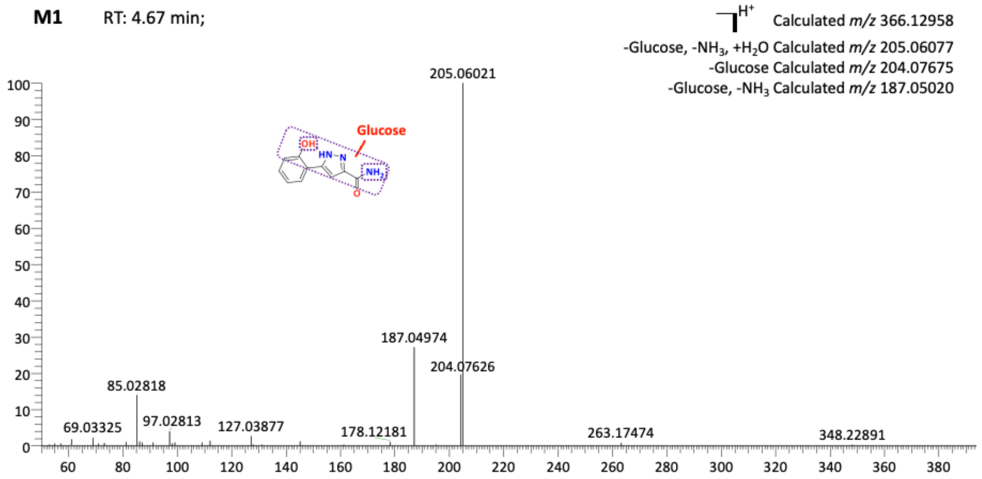


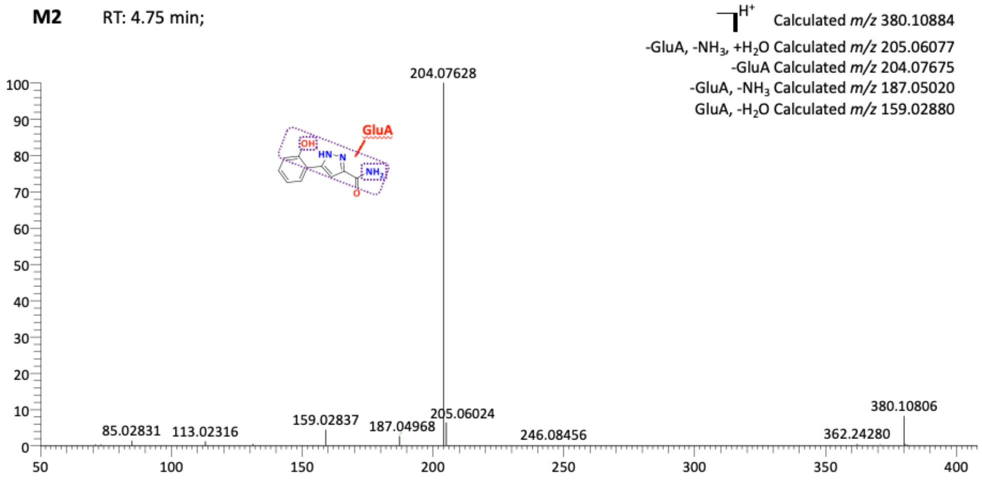


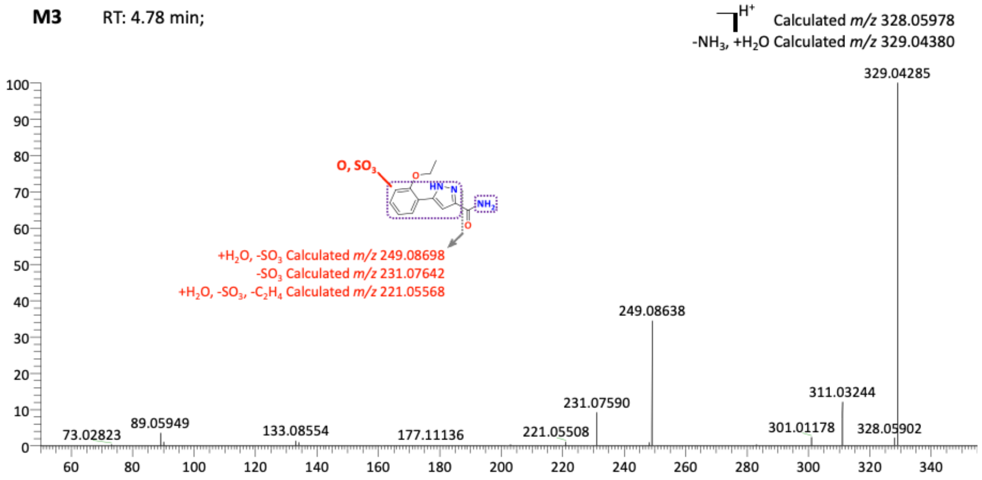

**M4** RT: 5.54 min;

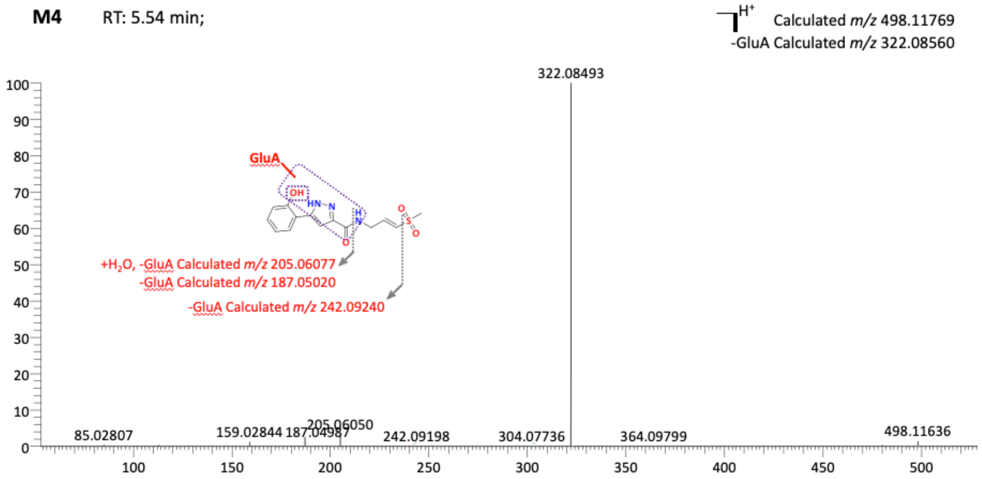


**M5** RT: 5.60 min;

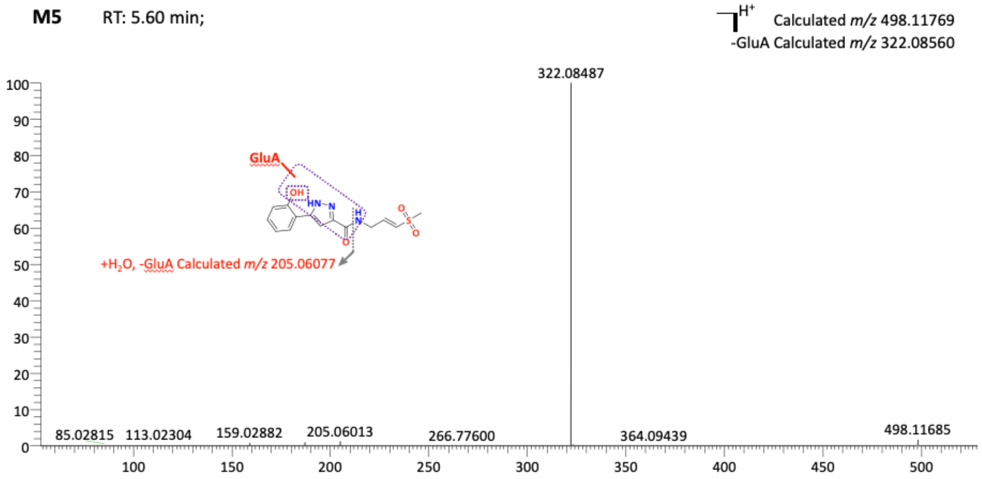


**M6** RT: 5.61 min;

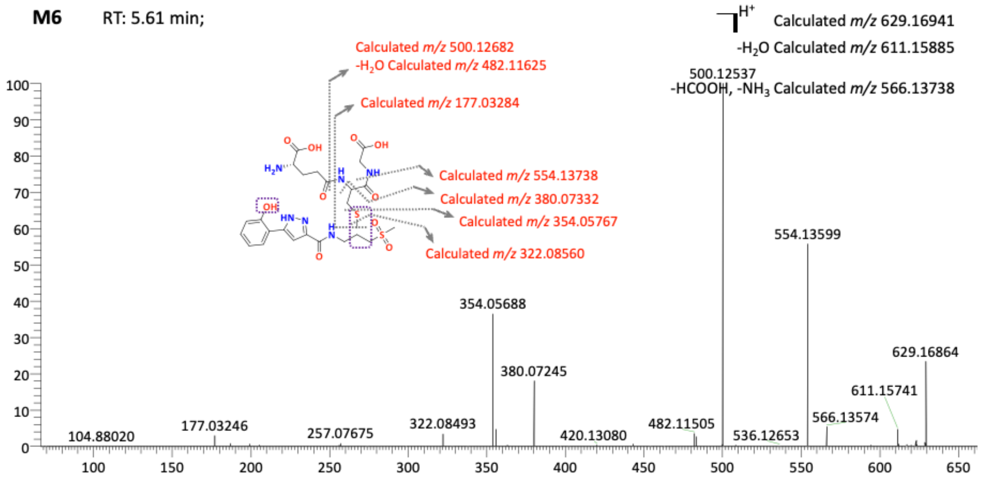

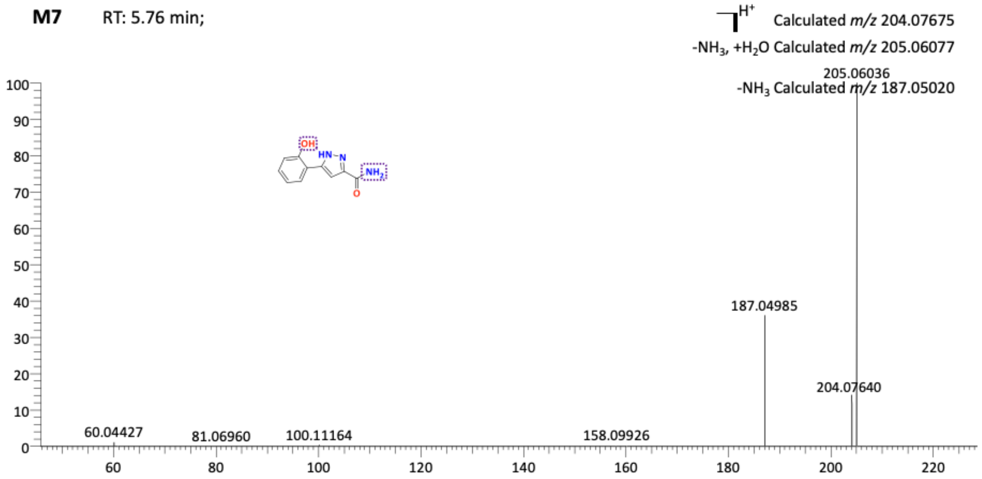
M7
RT: 5.76 min;
H+ Calculated m/z 204.07675
-NH3, +H2O Calculated m/z 205.06077
-NH3 Calculated m/z 187.05020
205.06036
187.04985
204.07640
60.04427
81.06960
100.11164
158.09926

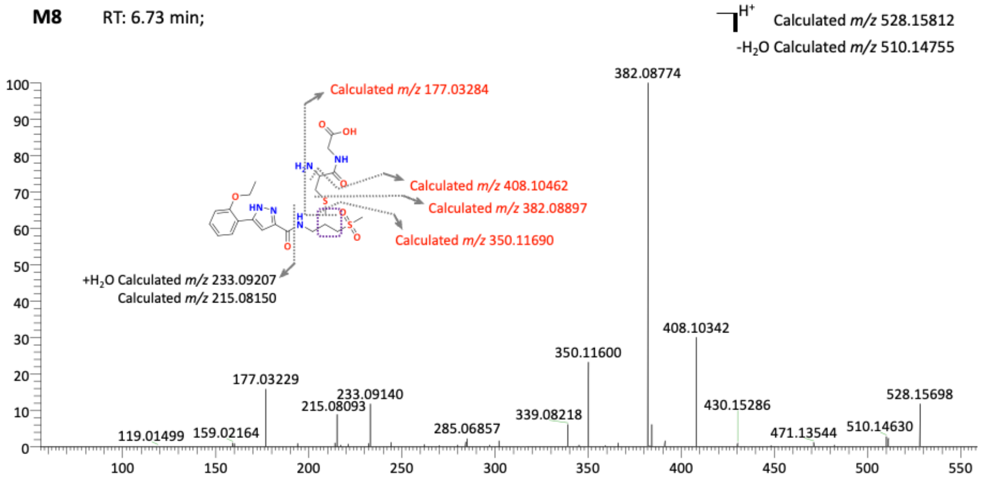
M8
RT: 6.73 min;
H+ Calculated m/z 528.15812
-H2O Calculated m/z 510.14755
Calculated m/z 177.03284
Calculated m/z 408.10462
Calculated m/z 382.08897
Calculated m/z 350.11690
+H2O Calculated m/z 233.09207
Calculated m/z 215.08150
382.08774
408.10342
350.11600
177.03229
233.09140
215.08093
285.06857
339.08218
430.15286
471.13544
510.14630
528.15698
119.01499
159.02164

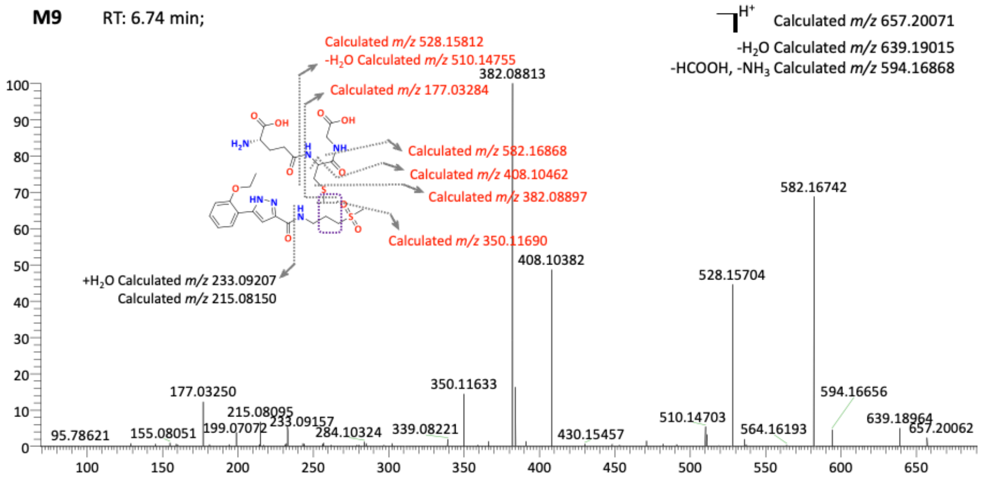
M9
RT: 6.74 min;
H+ Calculated m/z 657.20071
-H2O Calculated m/z 639.19015
-HCOOH, -NH3 Calculated m/z 594.16868
Calculated m/z 528.15812
-H2O Calculated m/z 510.14755
Calculated m/z 177.03284
Calculated m/z 582.16868
Calculated m/z 408.10462
Calculated m/z 382.08897
Calculated m/z 350.11690
+H2O Calculated m/z 233.09207
Calculated m/z 215.08150
382.08813
582.16742
408.10382
528.15704
350.11633
177.03250
215.08095
199.07072
233.09157
284.10324
339.08221
95.78621
155.08051
430.15457
510.14703
564.16193
594.16656
639.18964
657.20062

**M10** RT: 6.76 min;

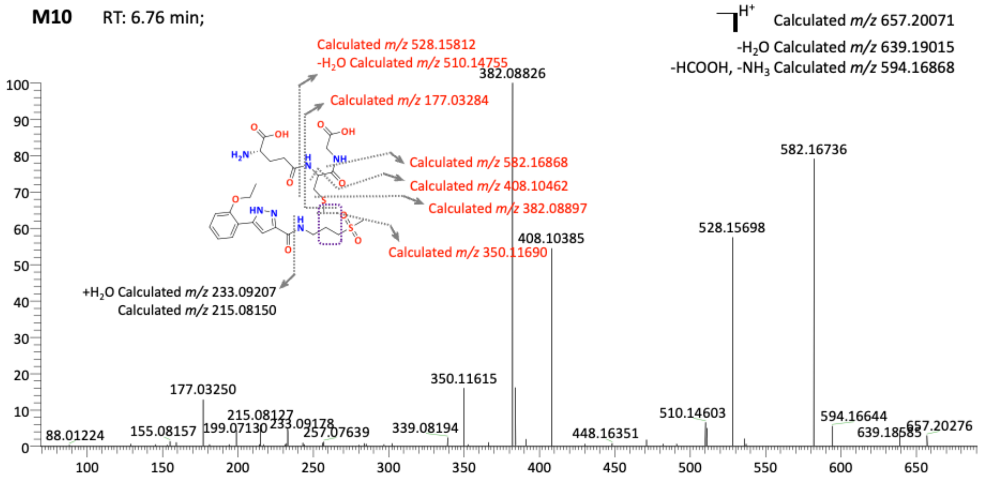


**M11** RT: 6.79 min;

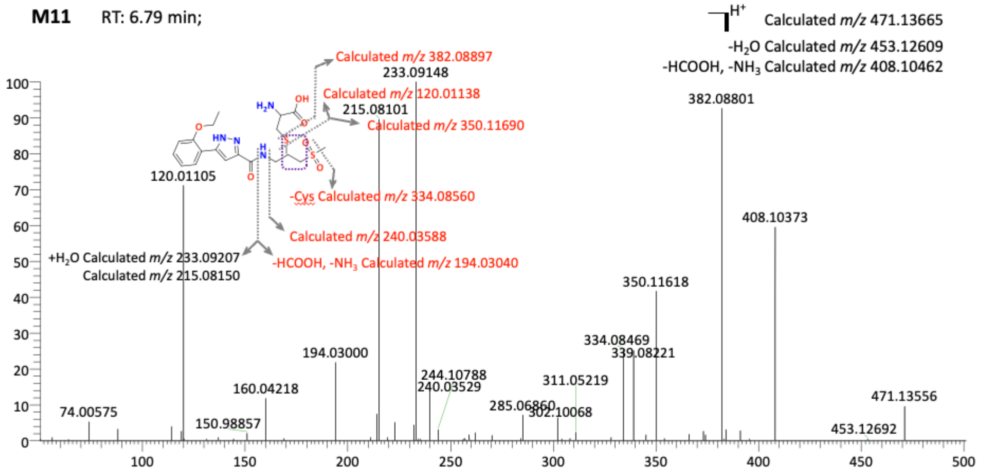


**M12** RT: 6.97 min;

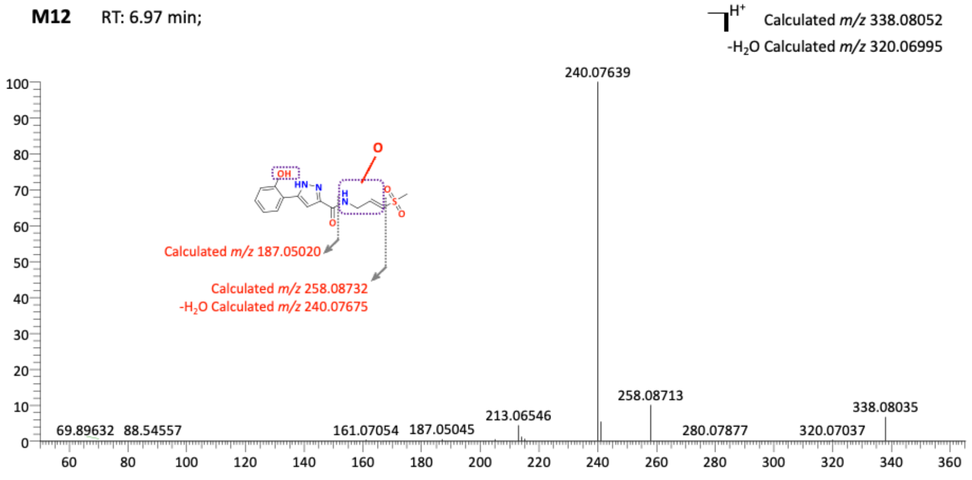

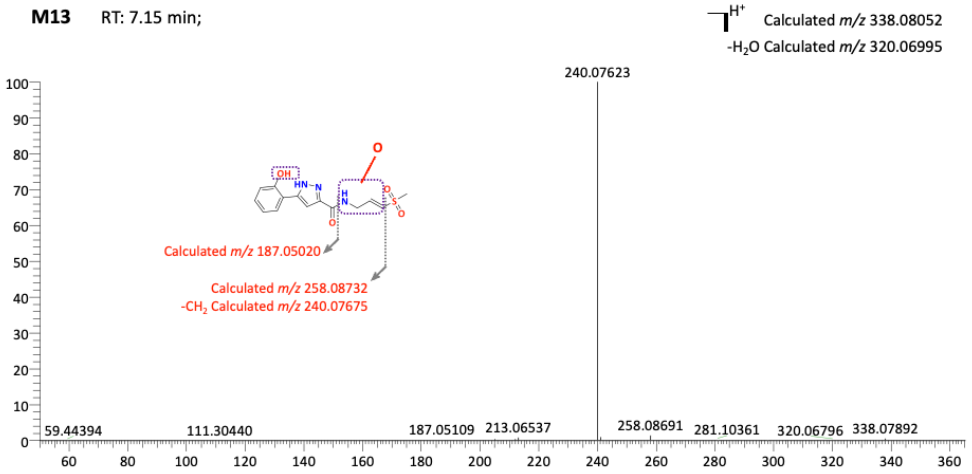
M13 RT: 7.15 min;
Calculated m/z 338.08052
-H2O Calculated m/z 320.06995
240.07623
Calculated m/z 187.05020
Calculated m/z 258.08732
-CH2 Calculated m/z 240.07675
59.44394
111.30440
187.05109
213.06537
258.08691
281.10361
320.06796
338.07892

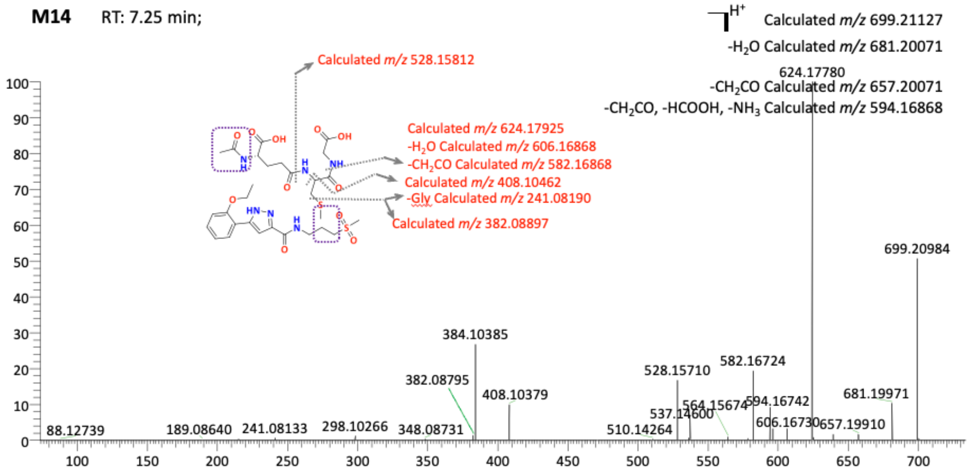
M14 RT: 7.25 min;
Calculated m/z 699.21127
-H2O Calculated m/z 681.20071
-CH2CO Calculated m/z 657.20071
-CH2CO, -HCOOH, -NH3 Calculated m/z 594.16868
Calculated m/z 528.15812
Calculated m/z 624.17925
-H2O Calculated m/z 606.16868
-CH2CO Calculated m/z 582.16868
Calculated m/z 408.10462
-Gly Calculated m/z 241.08190
Calculated m/z 382.08897
624.17780
699.20984
384.10385
382.08795
408.10379
528.15710
582.16724
681.19971
88.12739
189.08640
241.08133
298.10266
348.08731
510.14264
537.14600
564.15674
594.16742
606.16730
657.19910

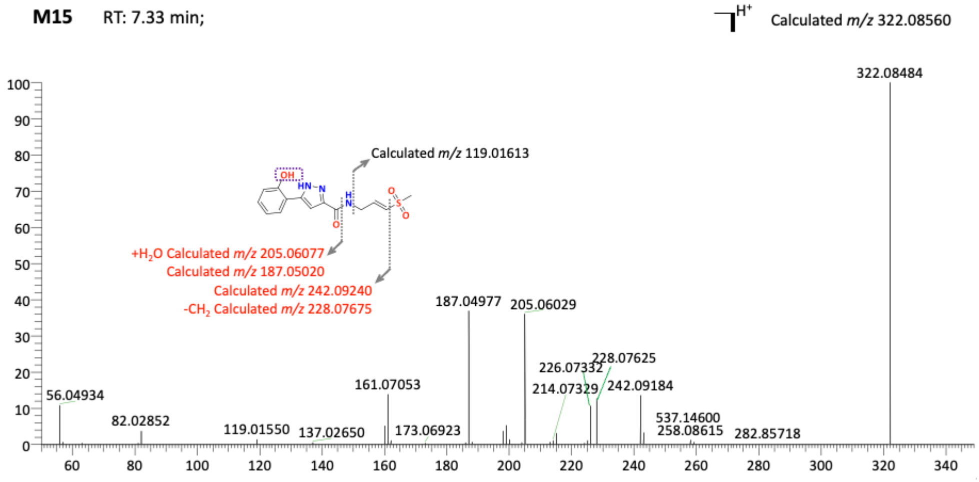
M15 RT: 7.33 min;
Calculated m/z 322.08560
322.08484
Calculated m/z 119.01613
+H2O Calculated m/z 205.06077
Calculated m/z 187.05020
Calculated m/z 242.09240
-CH2 Calculated m/z 228.07675
187.04977
205.06029
226.07332
228.07625
214.07329
242.09184
161.07053
56.04934
82.02852
119.01550
137.02650
173.06923
537.14600
258.08615
282.85718

**M16** RT: 7.52 min;

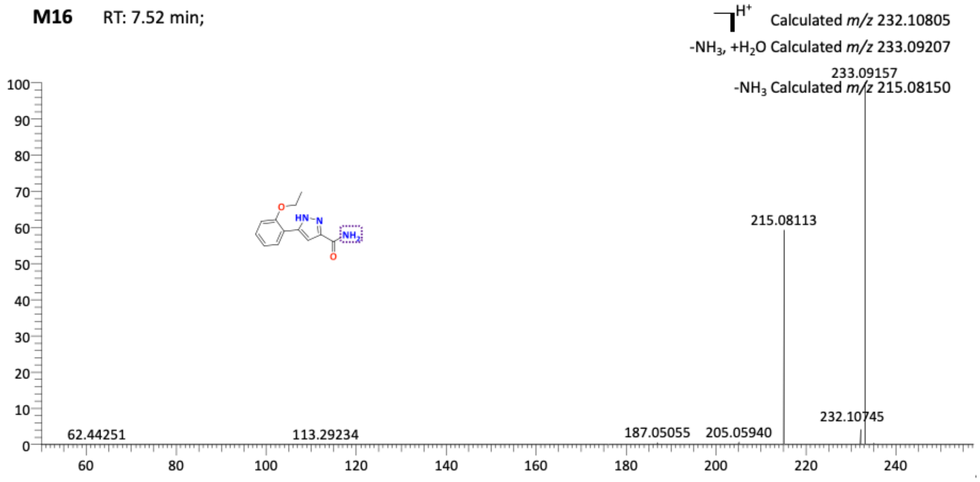


**M17** RT: 7.79 min;

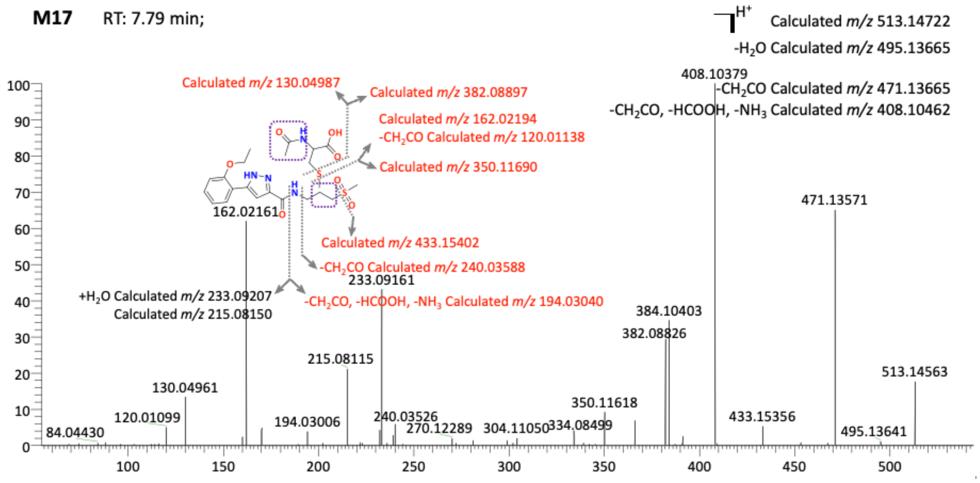


**M18** RT: 7.98 min;

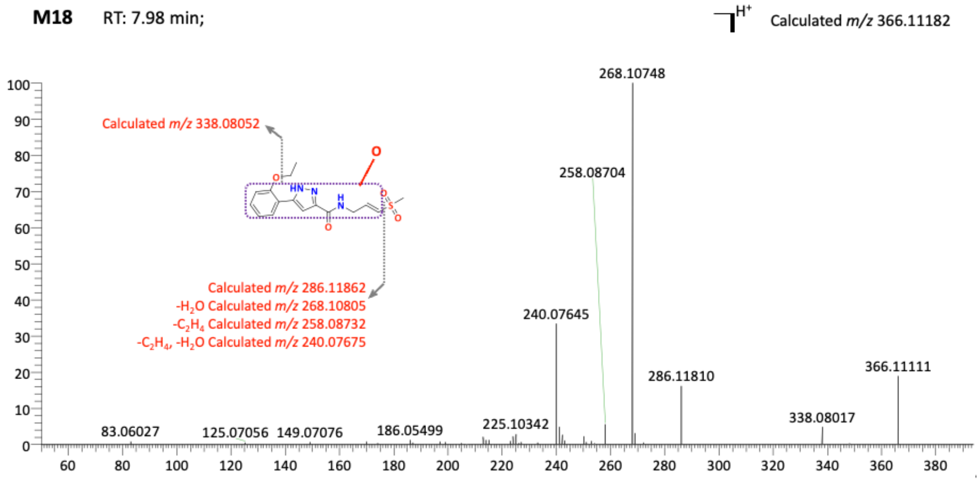

**M19** RT: 8.03 min;

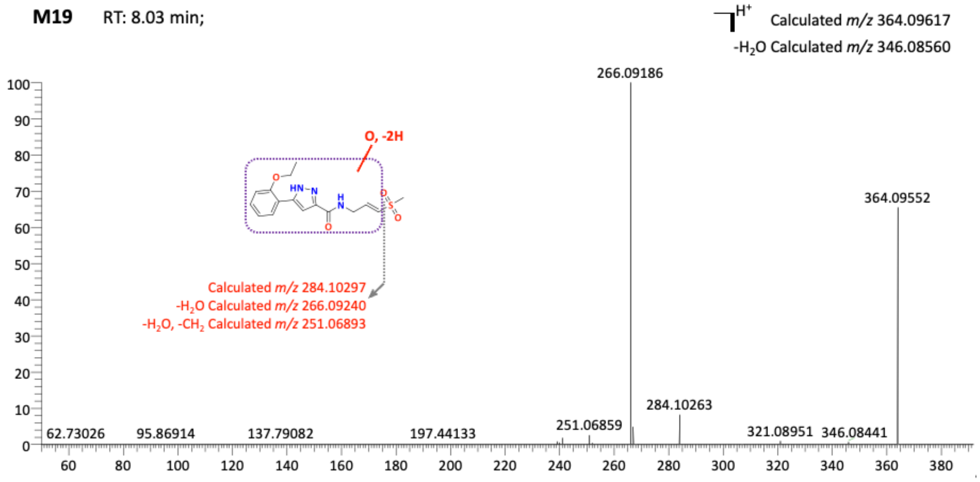


**M20** RT: 8.08 min;

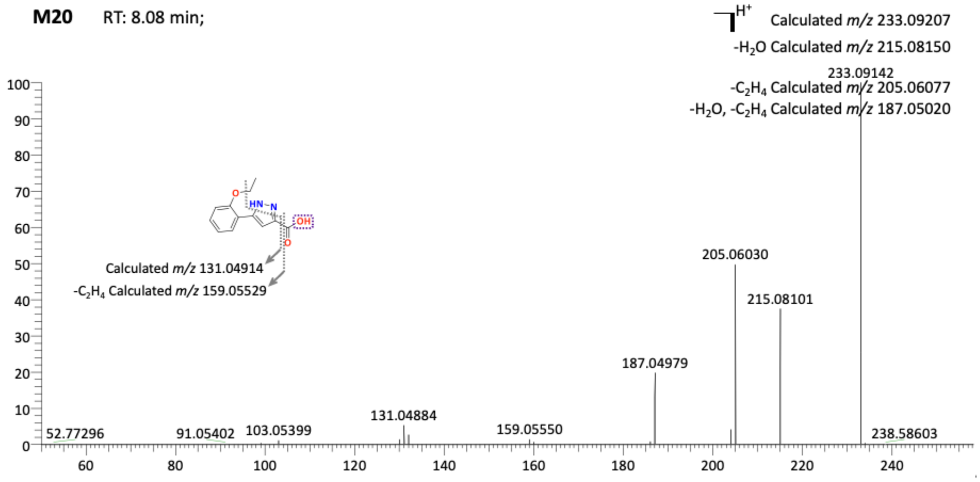


**M21** RT: 8.39 min;

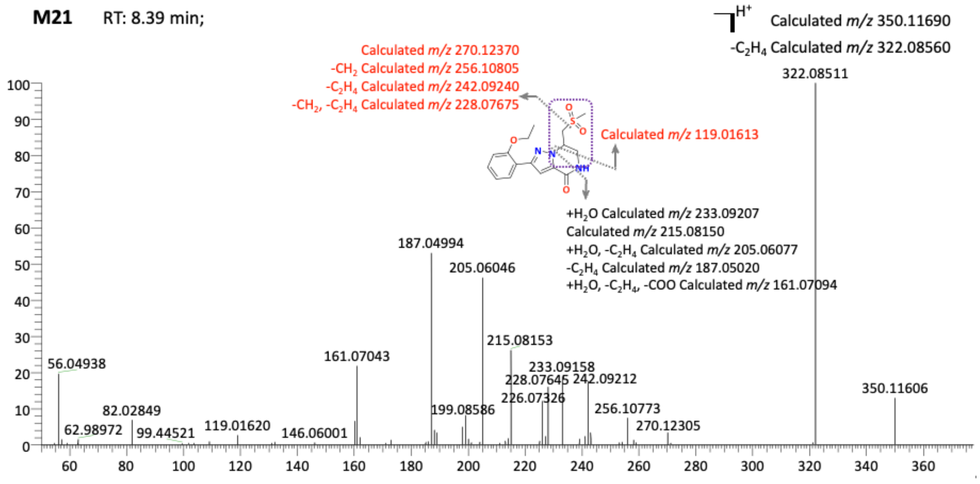

**M22** RT: 8.50 min;

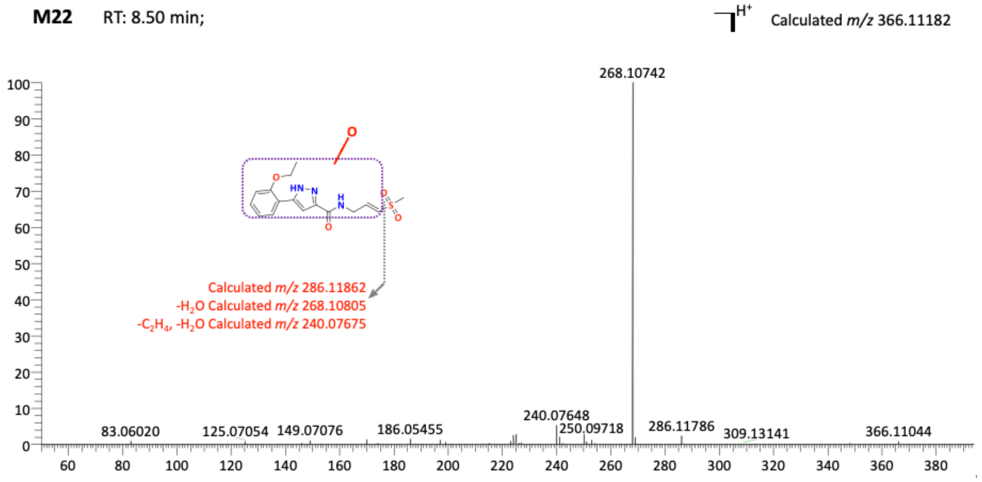
